# Non-collagenous and collagenous peptide markers for distinguishing between closely-related species using LC-MS/MS: A case study in baleen whales

**DOI:** 10.64898/2026.11.18.767015

**Authors:** Mitra Mohammadi Bazargani, Danielle L. Buss, Jean-Simon Richard, Mohsen Falahati-Anbaran, Brenna A. Frasier, Eirik Sollid, Lars Hagen, Cristina Brito, James H. Barrett

## Abstract

1. Reliable taxonomic identification of fragmented bone remains is central to ecological, evolutionary, archaeological and paleoecological research, yet existing proteomic approaches such as peptide mass fingerprinting (ZooMS) are sometimes constrained by peptide conservation in collagen, limiting resolution to the family-level. Moreover, ancient and curated samples are often small, and degraded, limiting the feasibility of techniques that require substantial amounts of and/or well-preserved starting material to ensure taxonomic identification.
2. Here, we develop and validate a minimally destructive Liquid Chromatography-Tandem Mass Spectrometry (LC-MS/MS) workflow that leverages both collagenous and non-collagenous bone proteins from small sample sizes (c. 0.1 mg of material) to improve genus and species-level discrimination in closely related taxa.
3. Using two morphologically similar and historically exploited whale species, the bowhead whale (*Balaena mysticetus*) and the North Atlantic right whale (*Eubalaena glacialis*), as a test case, we demonstrate that targeted sequencing of non-collagenous proteins, specifically prothrombin, yields consistent and diagnostic peptide markers that reliably distinguish between taxa that cannot be separated using ZooMS alone. We further evaluate recently proposed collagen peptide markers generated via LC-MS/MS and demonstrate that their discriminatory power is sometimes limited due to shared single amino-acid polymorphisms between taxa, resulting in overlapping peptide masses and indistinguishable amino-acid sequences for some individuals of both species.
4. Our pipeline requires c.0.1 mg of prepared sample, performs reliably on degraded material, and provides a cost-effective alternative when other techniques (e.g. ancient DNA) are not feasible.
5. By evaluating collagenous and non-collagenous protein markers, this framework increases the reliability of taxonomic identification while minimising sample destruction, and is applicable across ecological, evolutionary and heritage contexts requiring accurate identification from limited starting material.

## 1 Introduction

Proteomics has become an essential tool for taxonomic identification of fragmentary and morphologically ambiguous biological materials, particularly in archaeological, palaeontological and museum contexts (Coutu et al., 2021; Gilbert et al., 2024; Janzen et al., 2021). In studies focused on bone-based taxonomic identification, peptide mass fingerprinting, commonly termed Zooarchaeology by Mass Spectrometry (ZooMS), has emerged as the dominant approach (Buckley et al., 2009; van den Hurk et al., 2023; Warinner et al., 2022). ZooMS relies on enzymatic digestion of bone collagen followed by measurement of the resulting peptide mass fragments, typically using Matrix-Assisted Laser Desorption/Ionization Time-of-Flight Mass Spectrometry (MALDI-TOF MS), and comparing them to reference databases of known peptide mass profiles to assign taxonomic identity (e.g. Buckley et al., 2009; van den Hurk et al., 2023). This approach has enabled large-scale identification of faunal assemblages (Richter et al., 2022) and has substantially advanced reconstructions of past biodiversity, species distributions and human resource use (Buss et al., 2025; van den Hurk et al., 2023).

Despite its utility, ZooMS is intrinsically limited by the evolutionary conservation of fibrillar collagen. The structural constraints required to maintain collagen’s triple-helical architecture impose strong selection, resulting in limited sequence divergence among closely related taxa (Welker et al., 2015). Consequently, collagen peptide mass fingerprints are sometimes indistinguishable at the species or genus level, restricting taxonomic resolution in systems involving recently diverged or morphologically similar lineages (Richter et al., 2022).

Liquid Chromatography-Tandem Mass Spectrometry (LC-MS/MS) provides higher analytical resolution by sequencing peptides rather than relying solely on cleaved-peptide mass profiles. In this approach, enzymatic digestion of proteins producing peptides is followed by liquid chromatography, whereby peptides are separated by size and subsequently analysed by tandem mass spectrometry, in which selected precursor ions are fragmented and the resulting b- and y-ion series are measured. Because each amino acid has a characteristic monoisotopic mass, the mass differences between sequential fragment ions correspond to individual residues, enabling reconstruction of amino acid sequences and detection of single substitutions (with the exception of the isobaric residues leucine and isoleucine that are indistinguishable) (Steen & Mann, 2004). Although LC-MS/MS has been applied in palaeoproteomics and archaeological science, species identification workflows remain predominantly collagen-focused due to collagen’s abundance and long-term preservation in bone (Engels et al., 2025).

Non-collagenous proteins (NCPs), although less abundant than fibrillar collagen, are subject to fewer structural constraints and therefore often retain greater interspecific sequence variability (Wadsworth & Buckley, 2014). In contrast, collagen evolves relatively slowly, and collagen mass fingerprints are sometimes unable to discriminate among recently diverged or morphologically similar taxa, limiting taxonomic resolution in some groups, including cetaceans and pinnipeds (Buckley et al., 2009; 2014). Several studies have shown that archaeological bone preserves a diverse range of endogenous NCPs, including alpha-2-HS-glycoprotein (A2HSG/Fetuin-A), biglycan, chondroadherin, pigment epithelium-derived factor (PEDF), lumican, and prothrombin (Procopio, Mein, et al., 2021; Ntasi et al., 2022; Sawafuji et al., 2017; Wadsworth & Buckley, 2014), with potential applications in species identification and phylogenetic analysis (Giuffrida et al., 2018; Lanigan et al., 2020). Prothrombin, in particular, has been repeatedly identified in archaeological bone proteomes, suggesting strong preservation potential (Procopio, Hopkins, et al., 2021; Wadsworth & Buckley, 2014, 2018). Although Prothrombin has not previously been developed as a dedicated taxonomic marker protein, its consistent recovery and the presence of interspecific peptide variation make it a promising candidate for comparative taxonomic analyses.

Despite increasing recognition of the taxonomic potential and preservation rates of NCPs, few studies have directly evaluated their discriminatory power relative to collagen, especially in marine mammal archaeology (Buckley et al., 2014). While recent palaeoproteomic studies have identified whale-derived NCPs in archaeological remains, these proteins have generally been used opportunistically for tissue confirmation rather than as components of validated taxonomic marker systems. For instance, the muscle proteins and baleen keratins found in 16th-century whale remains were used to confirm bowhead whale tissue, but they were not applied as reusable markers for comparative species identification (Solazzo & Fitzhugh, 2024). Consequently, the utility of NCPs as reproducible taxonomic markers in marine mammal archaeology remains largely untested.

Recent studies have demonstrated recovery of NCPs from deep-time contexts, including the identification of serum albumin, fetuin-A, biglycan, chondroadherin, PEDF, lumican, and prothrombin in Early–Middle Pleistocene bone (Wadsworth & Buckley, 2014) and the persistence of diverse NCPs even in poorly preserved archaeological material (Procopio, Hopkins, et al., 2021). However, systematic evaluation of NCPs as diagnostic markers for genus and species-level identification of archaeological and paleontological fauna remains limited.

Alternatively, ancient DNA (aDNA) sequencing can provide high-resolution taxonomic and population-level information, yet aDNA preservation is highly variable, laboratory procedures are resource intensive, and destructive sampling requirements may preclude its use in small or culturally sensitive specimens (Gaeta, 2021; Pääbo et al., 2004; Shapiro & Hofreiter, 2014). Proteomic approaches can operate on minimal quantities of material (c. 0.1mg), are less susceptible to contamination concerns, and offer a minimally destructive alternative when DNA-based methods are impractical; this is particularly essential in studies looking at subfossil specimens, whereby DNA is sometimes too degraded for current aDNA protocols.

Here, we test whether incorporation of non-collagenous proteins into LC-MS/MS workflows facilitates taxonomic discrimination in closely related taxa that are indistinguishable using collagen-based peptide mass fingerprinting alone. We use two sister taxa within Balaenidae that occupy the North Atlantic Ocean, the bowhead whale (*Balaena mysticetus*) and the North Atlantic right whale (*Eubalaena glacialis*), as a validation system because their collagen peptide profiles are highly conserved and thus indistinguishable using ZooMS. Specifically, we (i) identify and validate diagnostic non-collagenous peptide markers, (ii) assess the discriminatory performance of recently proposed collagen markers, and (iii) compare outputs across two commonly used proteomic search engines. Finally, using historical and prehistorical bone material from natural history and archaeological contexts, (hereafter collectively termed historical specimens, irrespective of date and find context) as an empirical test case, we (iv) demonstrate that combined LC-MS/MS analysis of collagenous and non-collagenous proteins achieves genus or species-level resolution while maintaining minimal sample destruction, and can shed light on past human-wildlife interactions.

## 2 Materials and Methods

### 2.1 Sample selection

To assess whether incorporation of non-collagenous proteins (NCPs) into LC–MS/MS workflows facilitates taxonomic discrimination, and to evaluate the reliability of recently proposed collagen markers (Dekker et al., 2024), we analysed 27 Balaenidae specimens comprising 13 controls of known taxonomic identity and 14 unidentified historical specimens previously assigned to family level (Balaenidae) using peptide mass fingerprinting (Buss et al., under review). Control material was used for marker discovery and validation, including seven bowhead whale (*Balaena mysticetus*) specimens (one modern and six historical) and six North Atlantic right whale (*Eubalaena glacialis*) specimens (three modern and three historical). Controls were used to assess discriminatory performance of collagen and non-collagenous markers. Species identities of control specimens had been established previously using aDNA (McLeod et al., 2008; Sigfúsdóttir 2024), morphological assessment (Buss et al., under review) or museum archival documentation regarding modern deaths (see Supplementary Table S1 for pre-existing taxonomic information). The 14 historical Balaenidae specimens of unknown genus or species were included as an applied case study to demonstrate the utility of the validated marker framework for resolving genus and species-level identifications in historical bone material.

### 2.2 Preparation of samples

Proteins were extracted from bone material following the protocol from (Buss et al., under review). For one sample, FOC3489, extraction followed the protocol detailed in (van den Hurk et al., 2023). Approximately 0.1 mg aliquots were subsampled from “collagen” extracts (which contain both collagenous and non-collagenous proteins). The subsamples were resuspended in 1000 µL of 50 mM ammonium bicarbonate (AmBic) solution, pH 8, following Buss (2021). Protein samples were reduced by adding 500 mM dithiothreitol (DTT) to a final concentration of 20 mM and mixing briefly. Reduced protein samples were incubated at 60 °C for 1 hour. Following reduction, alkylation was performed by adding 1 M iodoacetamide (IAA) to a final concentration of 40 mM with brief mixing. The reaction mixture was incubated at room temperature for 30 minutes in the dark. The alkylation reaction was quenched by adding 500 mM DTT to a final concentration of 10 mM (Thermo Fisher Scientific 2022). Samples were then digested by adding 10 µL of 0.4 µg/µL trypsin solution and incubating at 37 °C overnight. Following incubation, the samples were centrifuged at 14,000 rpm at room temperature for 1 minute. To stop the trypsin digestion, 1 µL of 0.1% (v/v) trifluoroacetic acid (TFA) was added to each sample (Buss, 2021; van den Hurk et al., 2023). Following digestion, samples were desalted using C18 tips conditioned with 0.1% TFA + 50% acetonitrile (ACN) and equilibrated with 0.1% TFA in water. The material was aspirated up to 100 µL and cycled through ten times (as starting volume was 1000 µL). After cycling the sample through the tip, the tip was washed with 0.1% TFA in water and eluted with 0.1% TFA + 50% ACN (conditioning buffer) (Brown et al., 2020; Buss, 2021; van den Hurk et al., 2023). Finally, peptides were dried by speedvac.

### 2.3 LC-MS/MS analysis

LC-MS/MS was performed on a timsTOF Pro (Bruker Daltonics) connected to a nanoElute (Bruker Daltonics) HPLC at the Proteomics and and Modomics Experimental Core Facility of NTNU (PROMEC), Trondheim, Norway. Dried peptides were resuspended in 40 µL 0.1% formic acid (FA) before analysis. Peptide separation was done using a Pepsep 25 (150 µm*25 cm) column with running buffers A (0.1% FA) and B (0.1% FA in acetonitrile) with a gradient from 2% B to 40% B for 75min. The timsTOF instrument was operated in the DDA PASEF mode with 10 PASEF scans per acquisition cycle and accumulation and ramp times of 100 ms each. The ‘target value’ was set to 20,000 and dynamic exclusion was activated and set to 0.4 min. The quadrupole isolation width was set to 2 Th for m/z < 700 and 3 Th for m/z > 800.

### 2.4 Species identification using diagnostic peptide markers

Discrimination between two sister taxa, *Balaena mysticetus* and *Eubalaena glacialis* was achieved through the identification of diagnostic single amino-acid polymorphisms (SAPs) in collagenous and non-collagenous bone proteins. Candidate markers were identified in control specimens of known taxonomic identity and validated for application to historical specimens (Section 2.5). Discrimination was based on the *a priori* information (e.g. from ZooMS) that the specimens to be identified derived from the family Balaenidae. For *Eubalaena glacialis*, proteomic identification was to genus because no attempt was made to differentiate the North Atlantic Right whale from *Eubalaena australis* and *Eubalaena japonica*. However, the results can be interpreted at species level because only *Eubalaena glacialis* inhabits the North Atlantic. For *Balaena mysticetus* (within a monotypic genus), the proteomic identification was to species.

Raw LC–MS/MS data were searched against curated reference databases comprising collagen alpha-chain sequences from *Eubalaena glacialis* and *Balaena mysticetus.* Specifically, the databases included collagen alpha-2(I) sequences of North Atlantic right whale (*Eubalaena glacialis*) (NCBI Reference Sequence: XP_061054543.1), collagen alpha-1(I) sequences of North Atlantic right whale (*Eubalaena glacialis*) (NCBI Reference Sequence: XP_061030802.1), and collagen alpha-1(I) sequences of bowhead whale (*Balaena mysticetus*) (http://www.bowhead-whale.org/annotations/details/bmy_06723T0/; Keane et al., 2015). In addition, the reference proteome of common minke whale (*Balaenoptera acutorostrata*), a close relative with an available complete whale proteome at the time of analysis (Uniprot-Taxon ID 9767), was incorporated to facilitate assignment of both collagenous and non-collagenous proteins from LC–MS/MS data in the absence of complete Balaenidae proteomes (Table 1). FASTA entries corresponding to three previously reported collagen peptide markers containing SAPs distinguishing bowheads from whales of the genus *Eubalaena* (Dekker et al., 2024; Table 1) were also incorporated into the reference databases, including the relevant collagen alpha-2(I) peptide for bowhead whale. This step was particularly important because only a partial sequence for collagen alpha-2(I) was available for bowhead whale at the time of writing and it could not therefore be included in its entirety in the curated reference databases.

**Table 1.** Table of reference data used for assignments.

| **Database component** | **Taxon** | **Protein/Peptide(s) included** | **Purpose in analysis** | **Source** |
|---|---|---|---|---|
| Complete reference proteome | Common minke whale | Whole proteome | Assigning collagenous and non-collagenous proteins in the absence of complete balaenid proteomes | Uniprot taxonomic identifier 9767 |
| Collagen alpha-1(I) sequence | North Atlantic right whale | COL1A1 | Identifying species-diagnostic collagen SAPs | NCBI Reference Sequence: XP_061030802.1 |
| Collagen alpha-2(I) sequence | North Atlantic right whale | COL1A2 | Identifying species-diagnostic collagen SAPs | NCBI Reference Sequence: XP_061054543.1 |
| Collagen alpha-1(I) sequence | Bowhead whale | COL1A1 | Identifying species-diagnostic collagen SAPs | Keane et al., 2015; http://www.bowhead-whale.org/annotations/details/bmy_06723T0/ |
| COL1A1_918-957 | Right whale (*Eubalaena* spp.) | AGEVGPPGPPGPAGEKG**A**PGTDGPAGAPGSPGPQGIAGQR | Comparison with published peptide markers | Dekker et al., 2024 |
|  | Bowhead whale | AGEVGPPGPPGPAGEKG**S**PGTDGPAGAPGSPGPQGIAGQR | Comparison with published peptide markers | Dekker et al., 2024 |
| COL1A1_1170-1213 | Right whale (*Eubalaena* spp.) | GRTGDAGPAGPPGPPGPPGPPGPPS**V**GYDFSFLPQPPQEKQDGGR | Comparison with published peptide markers | Dekker et al., 2024 |
|  | Bowhead whale | GRTGDAGPAGPPGPPGPPGPPGPPS**A**GYDFSFLPQPPQEKQDGGR | Comparison with published peptide markers | Dekker et al., 2024 |
| COL1A2_262-280 | Right whale (*Eubalaena* spp.) | GSDGSVGPVGPAGPLGSAGPPGFPGAPGPKGELGPVGNPGP**P**GPAGSR | Comparison with published peptide markers | Dekker et al., 2024 |
|  | Bowhead whale | GSDGSVGPVGPAGPLGSAGPPGFPGAPGPKGELGPVGNPGP**S**GPAGSR | Comparison with published peptide markers | Dekker et al., 2024 |
| Bowhead whale protein/genome database | Bowhead whale | Protein and genomic resources | BLAST confirmation of candidate SAPs | Keane et al., 2015; http://www.bowhead-whale.org |
| Right whale genome assembly | North Atlantic right whale | Genome assembly | BLAST confirmation of candidate SAPs | NCBI Assembly: GCA_028564815.1 |
| Prothrombin FASTA database | North Atlantic right whale | IV**E**GRDAEIGLAPWQVMLFR | Validation of non-collagenous SAP markers | NCBI Reference Sequence: XP_061059283.1 |
|  | Bowhead whale | IV**D**GRDAEIGLAPWQVMLFR | Validation of non-collagenous SAP markers | Keane et al., 2015; http://www.bowhead-whale.org; (bmy_04116T0) |
| Serotransferrin-like FASTA database | North Atlantic right whale | GSVDEFE**K**CHLAR | Validation of non-collagenous SAP markers | NCBI Reference Sequence: XP_061050636.1 |
|  | Bowhead whale | GSVDEFE**R**CHLAR | Validation of non-collagenous SAP markers | NCBI GenBank: AAG32047.1 |

To identify and validate candidate SAP-containing peptide sequences, all raw proteomic data were interrogated using BLAST searches against publicly available bowhead whale and North Atlantic right whale genomic and protein resources, including the Bowhead Whale Genome resource at www.bowhead-whale.org (Keane et al., 2015) and a North Atlantic right whale genome assembly available through NCBI (GCA_028564815.1).

For non-collagenous peptides, a custom FASTA database was created containing bowhead whale and North Atlantic right whale prothrombin and serotransferrin-like protein sequences drawn from (for prothrombin) bmy_04116T0 available at http://www.bowhead-whale.org/ (Keane et al., 2015) and NCBI Reference Sequence XP_061059283.1 and (for serotransferrin-like) NCBI GenBank AAG32047.1 and NCBI Reference Sequence XP_061050636.1) (Table 1).

In addition, using BLAST, the NCP unconventional myosin-X was detected in North Atlantic Right Whale NCBI database resources, but it could not be identified in the Bowhead Whale Genome Resource (Keane et al., 2015) or NCBI bowhead whale references (taxid27602). Unconventional myosin-X is thus not presently considered further.

Searches of the curated databases for purposes of evaluating taxonomic identification were performed using MaxQuant (v2.4.14.0) (Tyanova et al., 2016) and pFind 3 (v3.2.1) (Chi et al., 2018). Searches assumed tryptic digestion with up to two missed cleavages permitted. For MaxQuant analyses, of all 27 samples considered, protease specificity was defined as trypsin/P, allowing cleavage C-terminal to lysine or arginine unless followed by proline. The minimum peptide length was set to seven amino acids. Carbamidomethylation of cysteine was specified as a fixed modification, while oxidation (M), hydroxyproline, deamidation (N/Q), and protein N-terminal acetylation were included as variable modifications, using a max PTM setting of 5. False discovery rates (FDR) were controlled at 1% at the levels of peptide-spectrum matches (PSMs), proteins, and modification sites using a target–decoy approach.

Equivalent parameters were applied in pFind analyses, which were conducted on a subset of 21 of the samples in the study. Trypsin was specified as the digesting enzyme with up to two missed cleavages permitted, and precursor and fragment mass tolerances were set to ±20 ppm. Carbamidomethylation (C) was treated as a fixed modification, while oxidation (M), oxidation (P), N-terminal acetylation, and deamidation (N/Q) were specified as variable modifications. Target–decoy searching was implemented with FDR controlled at 1% at both peptide and protein levels. Because leucine and isoleucine are isobaric and cannot be reliably distinguished using standard LC–MS/MS fragmentation methods, substitutions involving these residues were treated as ambiguous and reported as L/I.

To account for post-mortem protein degradation and incomplete enzymatic digestion characteristic of ancient bone proteomes (Fahrner et al., 2021; Warinner et al., 2022), selected samples exhibiting reduced peptide recovery under fully tryptic constraints were additionally analysed using semi-specific tryptic searches. Semi-specific digestion permits cleavage at one canonical tryptic site while allowing non-specific cleavage at the opposite peptide terminus, thereby improving recovery of partially degraded peptides (Fahrner et al., 2021).

Subsequently, candidate diagnostic peptides were defined as peptides that: (i) contained amino acid substitutions differentiating *Balaena mysticetus* and *Eubaelena glacialis* reference sequences for specimens known to belong to the family Balaenidae; (ii) were reproducibly detected in controls of one species; (iii) were absent from controls of the alternate species under identical analytical conditions; and (iv) were supported by fragment ion spectra confirming the position of the amino acid substitution. All control specimens were successfully identified to genus or species level based on the peptides studied.

For each peptide, presence/absence was tabulated across all specimens. Recovery rates for candidate genus or species-diagnostic peptide markers were assessed using modern and historical control specimens of known taxonomic identity (case study specimens were excluded from this validation step because their taxonomic identity was not assumed *a priori*). For each candidate marker, recovery was calculated as the proportion of control specimens in which the expected taxon-specific peptide variant was detected under the applied LC–MS/MS search conditions. Peptides yielding non-diagnostic balaenid variants, including spectra containing both variant forms, were not considered positive genus or species-level identifications for taxon-diagnostic recovery calculations. Marker performance was evaluated separately for *Balaena mysticetus* and *Eubalaena glacialis* controls to assess both peptide recovery and taxonomic specificity. Taxonomic specificity was assessed based on the absence of the diagnostic peptide in controls of the alternate taxon analysed under identical analytical conditions.

In addition to taxon-diagnostic validation, general marker recovery was assessed across the full sample set to evaluate peptide detection independently of taxonomic interpretation. Recovery rates depending on sample age (modern vs. historical) were also assessed with case study samples grouped with historical control samples due to their degraded nature. Each marker was scored as recovered when any peptide or variant call was detected, including genus or species-diagnostic calls, non-diagnostic balaenid-level calls, and calls containing both SAP variant forms. Entries recorded as “Not detected” were treated as non-recoveries. These recovery rates therefore describe marker detection success rather than genus or species-level diagnostic success.

Because sample numbers differed among groups, 95% confidence intervals for peptide marker recovery rates were calculated using the Wilson score method for binomial proportions (implemented using the binom package in R (Dorai-Raj & Cascone 2026)). Confidence intervals were calculated independently for the modern, historical, and overall datasets to quantify uncertainty associated with sample size. In addition, repeated random subsampling was used to assess the effect of sampling depth on recovery-rate estimates. Subsampling was performed independently for each marker and sample grouping (modern, historical) by repeatedly selecting a fixed number of specimens without replacement and recalculating the proportion of specimens in which the marker was detected. Subsampling was initially performed at a depth of ten specimens for the overall and historical datasets. Additionally, recovery rates were calculated at a subsampling depth of four specimens to allow comparison among modern, historical, and overall datasets, corresponding to the number of modern specimens available. For each sampling depth, 1,000 random subsamples were generated, and the mean recovery rate and central 95% subsampling ranges were calculated.

To assess whether increasing specimen numbers was likely to increase the recovery of distinct peptide variants, sample-based rarefaction was used to examine variant accumulation across the overall and historical datasets. Each unique variant at a given peptide locus was treated separately, with specimens containing both forms scored as positive for both. Variant accumulation curves were generated using 10,000 random permutations using the vegan package in R (Oksanen et al., 2026). Mean variant accumulation and 95% rarefaction ranges were calculated at each sampling depth to assess whether additional sampling was likely to recover previously undetected peptide variants.

To facilitate application of validated markers to historical specimens, candidate peptides were additionally weighted according to their observed recovery and taxonomic specificity in the control dataset. Peptides exhibiting consistently high recovery and taxonomic specificity across modern and historical controls were considered higher-confidence markers for taxonomic assignment, whereas markers with lower recovery or reduced discriminatory resolution were treated with greater caution. The weighting framework and justifications can be seen in Supplementary Table S2. Peptide markers that distinguished between North Atlantic right whale and Bowhead were scored highest if not known to overlap with other Mysticete. This weighting framework was subsequently applied to unidentified Balaenidae specimens from the Magdalen Islands, Gulf of St Lawrence, Canada, as a case study.

### 2.5 Magdalen Islands case study: Historical Balaenidae specimens from the Gulf of St Lawrence

A previous study identified 25 historical Balaenidae specimens from the Magdalen Islands (MI), located in the central Gulf of St Lawrence on the east coast of Canada (Buss et al., under review). Radiocarbon dating indicated a wide temporal distribution spanning approximately 4000 BCE to the mid-20th century, suggesting long-term presence of Balaenidae in the region. Of these 25 specimens previously identified as Balaenidae, 14 were selected at random for the present case study and their radiocarbon dates calibrated as discussed further below.

The Magdalen Islands were seasonally occupied by Basque whalers during the 16th and 17th centuries; however, the species targeted locally were not clearly documented. In contrast, historical material from Red Bay on the Strait of Belle Isle and other whaling stations on the northern Shore of the Gulf of St Lawrence demonstrates that bowhead whales (*Balaena mysticetus*) were the primary target of 16th- to 17th-century Basque whaling, rather than North Atlantic right whales (*Eubalaena glacialis*) (McLeod et al., 2008). Both taxa belong to the Balaenidae and are morphologically similar. Today, only North Atlantic right whales are regularly observed in the Gulf (Durette-Morin et al., 2022). Although none of the MI specimens derived from archaeological contexts (most were beach finds), they hold the potential to indicate whether Indigenous communities and/or early commercial whalers in the Magdalen Islands region would have interacted with bowhead whales, North Atlantic right whales, or both. Resolving species identity within this assemblage provides an applied context in which to evaluate the utility of discriminatory collagenous and non-collagenous proteomic markers. Distinguishing bowhead and North Atlantic right whales is especially important in this geographical region where the range of the two taxa may have overlapped in the past, and/or where *Eubalaena glacialis* may have replaced *Balaena mysticetus* through time (McLeod et al., 2008; Durette-Morin et al., 2022).

#### 2.5.1 LC-MS/MS analysis and species assignments of unidentified Balaenidae

The validated diagnostic peptides were applied to the 14 historical Balaenidae specimens from the Magdalen Islands. Species assignments were based on the combined presence, recovery weighting, and taxonomic specificity of validated diagnostic peptides. Greater interpretive weight was assigned to markers demonstrating high recovery rates and consistent species exclusivity within the control dataset. Species assignments required detection of at least one validated species-specific peptide at 1% false discovery rate supported by diagnostic fragment ions. Where possible, assignments were confirmed across analytical platforms.

Taxonomic confidence scores (Supplementary Table S2) were assigned using the weighted multi-locus framework introduced in section 2.4. COL1A1 1170–1213 was assigned a weight of 3, being reproducibly recovered and found to be taxonomically specific, despite the fact that other collagen-derived markers show lower overall discriminatory certainty, probably due to the high evolutionary conservation of collagen sequences. Prothrombin and the collagen marker COL1A2 262–280 were assigned a weight of 2 because they were consistently recovered, but found to be taxonomically specific only if a specimen was previously known to belong to the family Balaenidae (through ZooMS, for example). For both prothrombin and COL1A2 262–280, NCIB BLASTP searches against whales (taxid.9721) indicated that *Eubalaena* variants were shared with species from other whale families (*Balaenoptera musculus, Balaenoptera acutorostrata, Eschrichtius robustus, Balaenoptera physalus, Megaptera novaeangliae* and *Balaenoptera ricei*), whereas there were no such matches for *Balaena*. The serotransferrin-like marker was assigned a weight of 1 because it was rarely recovered and also found to be taxonomically specific only if a specimen was previously known to belong to the family Balaenidae (e.g. through ZooMS). In this case, variants were shared with species from other whale families for both *Eubalaena* (*Balaenoptera acutorostrata*) and *Balaena* (*Balaenoptera musculus*, *Balaenoptera ricei*, *Eschrichtius robustus*, *Balaenoptera borealis* and *Balaenoptera physalus*). COL1A1 918–957 was assigned a weight of 0 because both peptide variants occurred within North Atlantic right whale specimens and therefore lacked relevant taxonomic specificity.

Specimen-level confidence scores were calculated as the cumulative sum of concordant validated marker weights. Confidence categories were defined as low (1–2), moderate (3–4), high (5–6), and very high (≥7). Confidence scores were intended as relative evidence metrics reflecting marker reproducibility, taxonomic specificity, and peptide recovery within individual specimens rather than formal probabilistic assignments.

#### 2.5.2 Radiocarbon dating and temporal distribution of historical Balaenidae specimens

Previously and newly generated radiocarbon dates were used to provide temporal context for the proteomic species assignments (Supplementary Table S1). Previous radiocarbon details for the Magdalen Islands material are described in Buss et al. (under review). For this study, radiocarbon dates were recalibrated using species-specific marine reservoir calibrations. To account for regional variation in the marine reservoir effect, we estimated a bowhead whale-specific ΔR value using known-age specimens from Red Bay, Labrador. Red Bay represents a historically constrained sixteenth- to early seventeenth-century Basque whaling context, with the chronology of the relevant deposits restricted to 1536–1632 CE (McLeod et al., 2008, Table 1). Five bowhead specimens from Red Bay were thus used to estimate a regional offset (i.e. ΔR) from Marine20, the Northern Hemisphere marine calibration curve (Heaton et al., 2020): FOC3800, FOC3802, FOC3803, FOC3804, and FOC3820. The five known-age radiocarbon determinations were modelled within a single phase in OxCal (Bronk Ramsey 2001; Bronk Ramsey 2009) and assigned a shared ΔR parameter. Individual specimens were constrained between 1536 ± 25 and 1632 ± 25 CE using calendar-date priors.

The model produced a ΔR estimate of 155 ± 35 radiocarbon years. This estimate was subsequently used together with Marine20 to reservoir-correct all Gulf of St. Lawrence bowhead whale radiocarbon determinations. The uncalibrated and calibrated radiocarbon dates are shown in Supplementary Table S1 together with the proteomic results.

Kernel density estimation (KDE) was used to visualise clustering in the calibrated date distribution. Ninety-five percent confidence intervals were generated via bootstrapping, and local maxima and minima were extracted using the ‘findpeaks’ function (pracma package (Borchers 2025)) based on a 5% threshold of maximum density. Analyses were conducted in R version 4.5.1, with KDE models also visualised in OxCal (Bronk Ramsey 2017).

## 3 Results

### 3.1 Data generation

Raw LC–MS/MS data were successfully generated for all 27 analysed specimens, including modern controls, historical controls, and historical case study samples previously identified only as Balaenidae. Protein recovery was sufficient for genus or species-level peptide identification across both modern and historical material despite variable preservation states among historical specimens (Supplementary Table S3). Semi-specific tryptic searches improved peptide recovery in some degraded samples, particularly historical specimens FOC4291 and FOC4238 (Table 2).

**Table 2.**
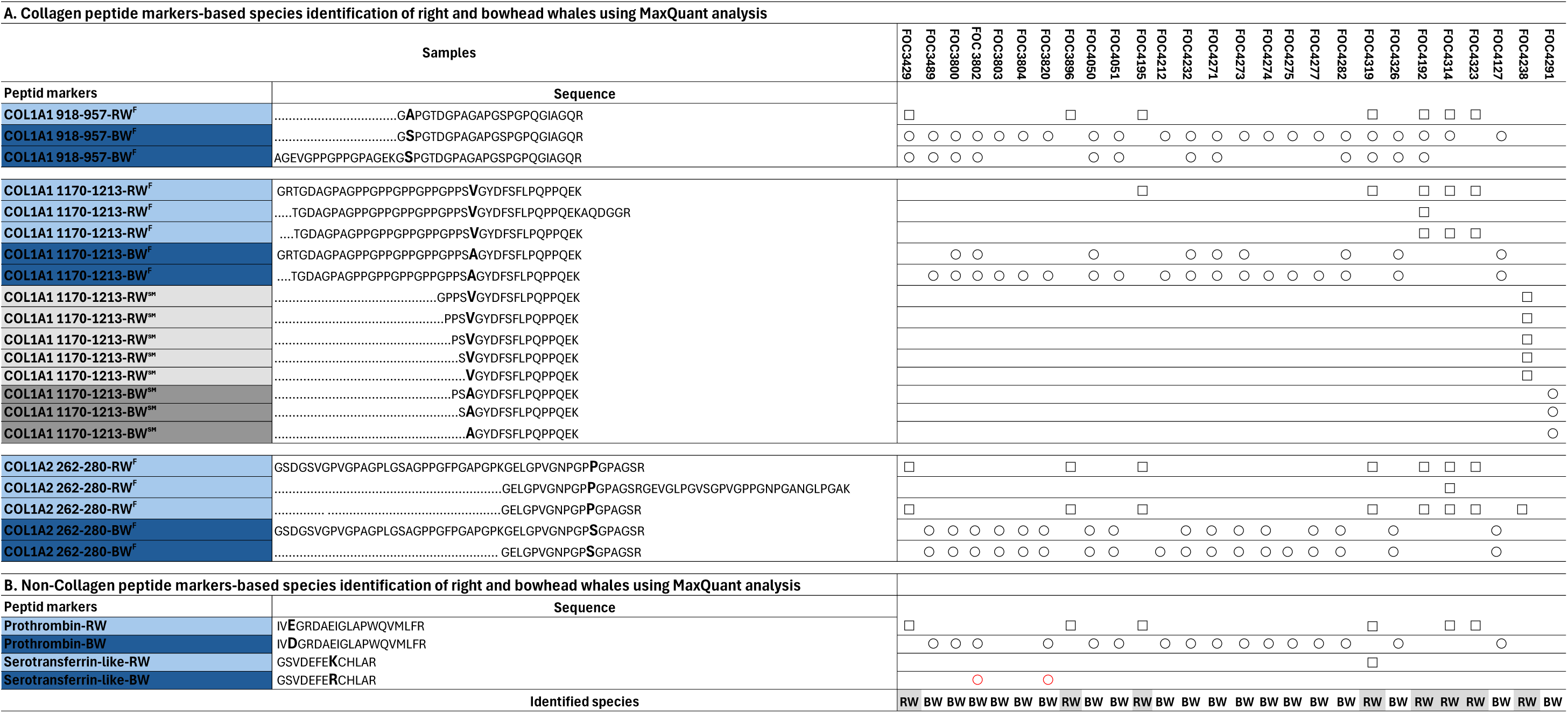
Peptide marker-based species identification of North Atlantic right whale and bowhead whale using MaxǪuant analysis. A. Collagen peptide markers, B. Non-collagenous peptide markers. □: identified by LC MS/MS as right whale. ○: identified by LC MS/MS as bowhead whale. FOC431S is a modern North Atlantic right whale and FOC432C a modern bowhead whale. The remaining samples are subfossil or historical. F: indicates trypsin/P (First analysis), Dark and light blue peptide markers were identified in the first analysis with full trypsin (trypsin/P). SM: indicates semi-trypsin, Dark and light gray peptide markers were identified in the second analysis with semi-trypsin.

| A. Collagen peptide markers-based species identification of right and bowhead whales using MaxQuant analysis |  |  |  |  |  |  |  |  |  |  |  |  |  |  |  |  |  |  |  |  |  |  |  |  |  |  |  |  |
| Samples |  | FOC3429 | FOC3489 | FOC3800 | FOC 3802 | FOC3803 | FOC3804 | FOC3820 | FOC3896 | FOC4050 | FOC4051 | FOC4195 | FOC4212 | FOC4232 | FOC4271 | FOC4273 | FOC4274 | FOC4275 | FOC4277 | FOC4282 | FOC4319 | FOC4326 | FOC4192 | FOC4314 | FOC4323 | FOC4127 | FOC4238 | FOC4291 |
| Peptid markers | Sequence |  |  |  |  |  |  |  |  |  |  |  |  |  |  |  |  |  |  |  |  |  |  |  |  |  |  |  |
|---|---|---|---|---|---|---|---|---|---|---|---|---|---|---|---|---|---|---|---|---|---|---|---|---|---|---|---|---|
| COL1A1 918-957-RW<sup>F</sup> | ..............................G**A**PGTDGPAGAPGSPGPQGIAGQR | □ |  |  |  |  |  |  | □ |  |  | □ |  |  |  |  |  |  |  |  | □ |  | □ | □ | □ |  |  |  |
| COL1A1 918-957-BW<sup>F</sup> | ..............................G**S**PGTDGPAGAPGSPGPQGIAGQR | ○ | ○ | ○ | ○ | ○ | ○ | ○ |  | ○ | ○ |  | ○ | ○ | ○ | ○ | ○ | ○ | ○ | ○ | ○ | ○ | ○ | ○ |  | ○ |  |  |
| COL1A1 918-957-BW<sup>F</sup> | AGEVGPPGPPGPAGEKG**S**PGTDGPAGAPGSPGPQGIAGQR | ○ | ○ | ○ | ○ |  |  |  |  | ○ | ○ |  |  | ○ | ○ |  |  |  |  | ○ | ○ | ○ | ○ |  |  |  |  |  |
| COL1A1 1170-1213-RW<sup>F</sup> | GRTGDAGPAGPPGPPGPPGPPGPPS**V**GYDFSFLPQPPQEK |  |  |  |  |  |  |  |  |  |  | □ |  |  |  |  |  |  |  |  | □ |  | □ | □ | □ |  |  |  |
| COL1A1 1170-1213-RW<sup>F</sup> | .....TGDAGPAGPPGPPGPPGPPGPPS**V**GYDFSFLPQPPQEKAQDGGR |  |  |  |  |  |  |  |  |  |  |  |  |  |  |  |  |  |  |  |  |  | □ |  |  |  |  |  |
| COL1A1 1170-1213-RW<sup>F</sup> | ....TGDAGPAGPPGPPGPPGPPGPPS**V**GYDFSFLPQPPQEK |  |  |  |  |  |  |  |  |  |  |  |  |  |  |  |  |  |  |  |  |  | □ | □ | □ |  |  |  |
| COL1A1 1170-1213-BW<sup>F</sup> | GRTGDAGPAGPPGPPGPPGPPGPPS**A**GYDFSFLPQPPQEK |  |  | ○ | ○ |  |  |  |  | ○ |  |  |  | ○ | ○ | ○ |  |  |  | ○ |  | ○ |  |  |  | ○ |  |  |
| COL1A1 1170-1213-BW<sup>F</sup> | ....TGDAGPAGPPGPPGPPGPPGPPS**A**GYDFSFLPQPPQEK |  | ○ | ○ | ○ | ○ | ○ | ○ |  | ○ | ○ |  | ○ | ○ | ○ | ○ | ○ | ○ | ○ | ○ |  | ○ |  |  |  | ○ |  |  |
| COL1A1 1170-1213-RW<sup>SM</sup> | ........................................GPPS**V**GYDFSFLPQPPQEK |  |  |  |  |  |  |  |  |  |  |  |  |  |  |  |  |  |  |  |  |  |  |  |  |  | □ |  |
| COL1A1 1170-1213-RW<sup>SM</sup> | ..........................................PPS**V**GYDFSFLPQPPQEK |  |  |  |  |  |  |  |  |  |  |  |  |  |  |  |  |  |  |  |  |  |  |  |  |  | □ |  |
| COL1A1 1170-1213-RW<sup>SM</sup> | ............................................PS**V**GYDFSFLPQPPQEK |  |  |  |  |  |  |  |  |  |  |  |  |  |  |  |  |  |  |  |  |  |  |  |  |  | □ |  |
| COL1A1 1170-1213-RW<sup>SM</sup> | .............................................S**V**GYDFSFLPQPPQEK |  |  |  |  |  |  |  |  |  |  |  |  |  |  |  |  |  |  |  |  |  |  |  |  |  | □ |  |
| COL1A1 1170-1213-RW<sup>SM</sup> | ...............................................**V**GYDFSFLPQPPQEK |  |  |  |  |  |  |  |  |  |  |  |  |  |  |  |  |  |  |  |  |  |  |  |  |  | □ |  |
| COL1A1 1170-1213-BW<sup>SM</sup> | ............................................PS**A**GYDFSFLPQPPQEK |  |  |  |  |  |  |  |  |  |  |  |  |  |  |  |  |  |  |  |  |  |  |  |  |  |  | ○ |
| COL1A1 1170-1213-BW<sup>SM</sup> | .............................................S**A**GYDFSFLPQPPQEK |  |  |  |  |  |  |  |  |  |  |  |  |  |  |  |  |  |  |  |  |  |  |  |  |  |  | ○ |
| COL1A1 1170-1213-BW<sup>SM</sup> | ...............................................**A**GYDFSFLPQPPQEK |  |  |  |  |  |  |  |  |  |  |  |  |  |  |  |  |  |  |  |  |  |  |  |  |  |  | ○ |
| COL1A2 262-280-RW<sup>F</sup> | GSDGSVGPVGPAGPLGSAGPPGFPGAPGPKGELGPVGNPGP**P**GPAGSR | □ |  |  |  |  |  |  | □ |  |  | □ |  |  |  |  |  |  |  |  | □ |  | □ | □ | □ |  |  |  |
| COL1A2 262-280-RW<sup>F</sup> | ........................................................GELGPVGNPGP**P**GPAGSRGEVGLPGVSGPVGPPGNPGANGLPGAK |  |  |  |  |  |  |  |  |  |  |  |  |  |  |  |  |  |  |  |  |  |  | □ |  |  |  |  |
| COL1A2 262-280-RW<sup>F</sup> | ............ ..........................................GELGPVGNPGP**P**GPAGSR | □ |  |  |  |  |  |  | □ |  |  | □ |  |  |  |  |  |  |  |  | □ |  | □ | □ | □ |  | □ |  |
| COL1A2 262-280-BW<sup>F</sup> | GSDGSVGPVGPAGPLGSAGPPGFPGAPGPKGELGPVGNPGP**S**GPAGSR |  | ○ | ○ | ○ | ○ | ○ | ○ |  | ○ | ○ |  |  | ○ | ○ | ○ | ○ |  | ○ | ○ |  | ○ |  |  |  | ○ |  |  |
| COL1A2 262-280-BW<sup>F</sup> | ....................................................... GELGPVGNPGP**S**GPAGSR |  | ○ | ○ | ○ | ○ | ○ | ○ |  | ○ | ○ |  | ○ | ○ | ○ | ○ | ○ | ○ | ○ | ○ |  | ○ |  |  |  | ○ |  |  |
| B. Non-Collagen peptide markers-based species identification of right and bowhead whales using MaxQuant analysis |  |  |  |  |  |  |  |  |  |  |  |  |  |  |  |  |  |  |  |  |  |  |  |  |  |  |  |  |
| Peptid markers | Sequence |  |  |  |  |  |  |  |  |  |  |  |  |  |  |  |  |  |  |  |  |  |  |  |  |  |  |  |
| Prothrombin-RW | IV**E**GRDAEIGLAPWQVMLFR | □ |  |  |  |  |  |  | □ |  |  | □ |  |  |  |  |  |  |  |  | □ |  |  | □ | □ |  |  |  |
| Prothrombin-BW | IV**D**GRDAEIGLAPWQVMLFR |  | ○ | ○ | ○ |  |  | ○ |  | ○ | ○ |  | ○ | ○ | ○ | ○ | ○ | ○ | ○ | ○ |  | ○ |  |  |  | ○ |  |  |
| Serotransferrin-like-RW | GSVDEFE**K**CHLAR |  |  |  |  |  |  |  |  |  |  |  |  |  |  |  |  |  |  |  | □ |  |  |  |  |  |  |  |
| Serotransferrin-like-BW | GSVDEFE**R**CHLAR |  |  |  | ○ |  |  | ○ |  |  |  |  |  |  |  |  |  |  |  |  |  |  |  |  |  |  |  |  |
|  | Identified species | RW | BW | BW | BW | BW | BW | BW | RW | BW | BW | RW | BW | BW | BW | BW | BW | BW | BW | BW | RW | BW | RW | RW | RW | BW | RW | BW |

### 3.2 Peptide Marker recovery

Five peptide markers were recovered across the dataset (COL1A1 918–957; COL1A1 1170–1213; COL1A2 262–280; prothrombin; serotransferrin-like). Peptide marker recovery varied substantially among the five candidate markers (Supplementary Table S4). Across all samples, the highest general recovery rate was observed for COL1A2 262–280 which was detected in 26/27 (96.3%; CI: 81.7–99.3%). COL1A1 1170–1213 and prothrombin were detected in 25/27 (92.6%; CI: 76.6–97.9%) and 22/27 (81.5%; CI: 63.3–91.8%) samples, respectively. COL1A1 918–957 was also frequently recovered, occurring in 25/27 samples (92.6%; CI: 76.6–97.9%), although this marker was not retained as genus or species-diagnostic because of polymorphisms that were shared across the Balaenidae taxa. The Serotransferrin-like marker was recovered less frequently (3/27, 11.1%; CI: 3.9–28.1%).

When only historical samples were considered, recovery rates slightly reduced in some markers. COL1A2 262–280 and COL1A1 1170-1213 remained the most consistently recovered genus and species-diagnostic markers, with recovery rates of 22/23 (95.7%; CI: 79.0–99.2%) and 21/23 (91.3%; 73.2–97.6%), respectively. Among non-collagenous markers, prothrombin was the most consistently recovered, appearing in 18/23 (78.3%; CI: 58.1–90.3%) of historical samples. In terms of modern samples, COL1A2 262–280, COL1A1 1170-1213 and prothrombin exhibited the highest recovery among genus and species-diagnostic markers, at 100%. For the overall and historical only datasets, repeated random subsampling at a depth of ten specimens produced recovery rates of 90–100% for COL1A2 262–280, 80–100% for COL1A1 1170–1213, and 60–100% for prothrombin in both datasets (see Table S4). A full list of recovery rates and results from accumulation and rarefaction analyses are available in Supplementary Tables S4 and S5.

Species-specific diagnostic recovery was assessed separately using known control specimens (Supplementary Table S5). COL1A2 262–280 showed the highest diagnostic recovery, with the expected species-specific variant identified in all 13 controls (100%). COL1A1 1170–1213 showed similarly high overall diagnostic recovery (11/13; 84.6%), although recovery differed between bowhead whale (7/7; 100%) and North Atlantic right whale controls (4/6; 66.7%). Prothrombin exhibited diagnostic recovery of 10/13 (76.9%), with the expected variant detected in 5/7 bowhead whale controls (71.4%) and 5/6 North Atlantic right whale controls (83.3%). Serotransferrin-like was recovered in 2/7 bowhead whale controls (28.6%) and 1/6 North Atlantic right whale controls (16.6%). In contrast, COL1A1 918–957 was excluded from diagnostic scoring because of overlapping polymorphisms between taxa.

Sample-based rarefaction suggested that peptide variant accumulation approached saturation at approximately 20 specimens, indicating that further sampling was unlikely to substantially increase the number of distinct variants recovered at the loci examined (Supplementary Figure S1).

### 3.3 Identification and validation of diagnostic peptide markers

Multiple collagenous and non-collagenous peptide markers containing SAPs capable of distinguishing bowhead whale and North Atlantic right whale were identified by MaxQuant analysis (Table 2).

Among the evaluated loci, a non-collagenous peptide derived from prothrombin showed strong and consistent discriminatory performance within the family Balaenidae. BLAST interrogation of bowhead whale and right whale genomic resources identified a glutamic acid/aspartic acid substitution at the third amino acid position of the peptide sequence (IVEGRDAEIGLAPWQVMLFR in North Atlantic right whale; IVDGRDAEIGLAPWQVMLFR in bowhead whale). Following incorporation of both peptide variants into a custom FASTA database, the prothrombin marker was detected in 22 of 27 analysed specimens, including all modern controls (4/4) and most historical specimens (18/23). Recovery of the bowhead-associated D amino acid substitution and North Atlantic right whale-associated E amino acid substitution was similarly high among all (both modern and historical) controls (5/7; 71.4% and 5/6; 83.3%, respectively). Where species-specific recovery was below 100%, this reflects failure to recover the relevant peptide marker in some controls rather than detection of the alternate taxon variant; when the prothrombin marker was recovered, the species-specific substitution was always correct. Marker recovery varied with sample condition, but diagnostic accuracy was 100% if the peptide was detected.

Three previously proposed collagen peptide markers from Dekker et al. (2024) were additionally evaluated. Of these, COL1A2 262–280 and COL1A1 1170–1213 showed high reproducibility and consistently differentiated the two taxa across both modern and historical material. For COL1A2 262–280, the bowhead-associated S amino acid substitution and North Atlantic right whale-associated P amino acid variant were recovered in all bowhead whale controls (7/7; 100%) and right whale controls (6/6; 100%) respectively. The COL1A1 1170–1213 marker with the bowhead-associated A amino acid substitution was detected in all bowhead whale controls (7/7; 100%) whereas the North Atlantic right whale-associated V amino acid substitution showed lower but still informative recovery in right whale controls (4/6; 66.7%). When assignments were not 100% it was because the peptide marker was not recovered, rather than it was recovered with the wrong marker. For the COL1A1 1170–1213 marker, semi-specific tryptic searches in MaxQuant improved recovery in selected degraded specimens (FOC4291 and FOC4238, for which the peptide was not detected under fully tryptic search conditions). It is notable that COL1A1 1170–1213 appears to be diagnostic for both *Balaena mysticetus and Eubalaena* without *a priori* knowledge that a specimen belongs to the family Balaenidae, unlike the other markers considered; a NCIB BLASTP search against whales (taxid.9721) did not yield matches from other taxonomic families (see Section 2.5.1).

The COL1A1 918–957 locus did not consistently discriminate between the two taxa. Although the bowhead-associated S amino acid substitution was detected in 100% (7/7) of bowhead whale controls, the same variant also occurred in multiple North Atlantic right whale controls, including both modern and historical specimens (Table 2). The North Atlantic right whale-associated amino acid substitution at this locus was exclusively recovered in only 33.3% (2/6) of right whale controls (Table 2). Several specimens yielded non-diagnostic balaenid peptide variants containing both amino acid substitution. Together, these observations indicate that COL1A1 918–957 lacks genus or species specificity and may produce erroneous assignments, especially in degraded ancient material where peptide drop-out limits marker recovery and where proline hydroxylation/oxidation may also be a complicating factor given that it produces a mass shift identical to an A to S substitution.

Lastly, the serotransferrin-like non-collagenous peptide was rarely detected, although it appears to be taxonomically specific if already known to be Balaenidae. The lysine/arginine (K) amino acid substitution was identified in 1 of 6 (16.6%) North Atlantic right whale controls and the R amino acid substitution was detected in 2 of 7 (28.6%) bowhead whale controls.

Overall, the prothrombin SAP and the collagen markers COL1A2 262–280 and COL1A1 1170–1213 provided the most reliable species discrimination across both modern and historical specimens and were therefore weighted most strongly in subsequent species assignments (see Section 2.5.1 for details). For a full list of marker recovery rates please see Supplementary Table S5.

### 3.4 Comparison of MaxQuant and pFind performance

Species assignments were broadly concordant between MaxQuant (Table 2) and pFind (Table 3) analyses. Both search engines consistently recovered validated diagnostic peptide markers, including the prothrombin locus, and supported identical specimen-level taxonomic assignments across all validated specimens that were analysed using both methods. Despite this concordance, the two search platforms differed in peptide recovery characteristics. MaxQuant showed stronger recovery of validated non-collagenous peptide markers, particularly prothrombin.

**Table 3.**
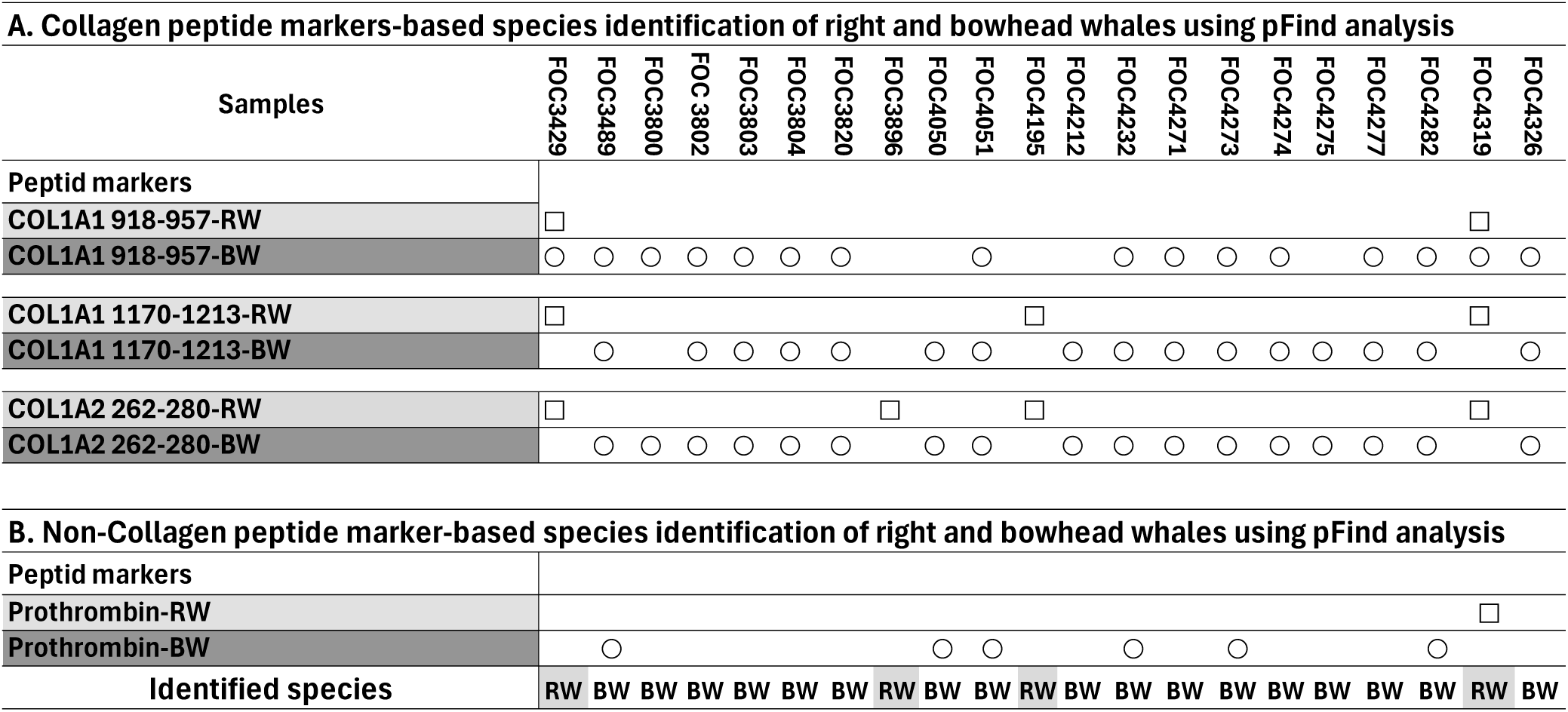
Peptide marker-based species identification of North Atlantic right whale and bowhead whale using pFind analysis. A. Collagen peptide markers, B. Non-collagenous peptide markers. *□: identified by LC MS/MS as right whale. ○: identified by LC MS/MS as bowhead whale*.

| A. Collagen peptide markers-based species identification of right and bowhead whales using pFind analysis |  |  |  |  |  |  |  |  |  |  |  |  |  |  |  |  |  |  |  |  |  |
|---|---|---|---|---|---|---|---|---|---|---|---|---|---|---|---|---|---|---|---|---|---|
| Samples | FOC3429 | FOC3489 | FOC3800 | FOC 3802 | FOC3803 | FOC3804 | FOC3820 | FOC3896 | FOC4050 | FOC4051 | FOC4195 | FOC4212 | FOC4232 | FOC4271 | FOC4273 | FOC4274 | FOC4275 | FOC4277 | FOC4282 | FOC4319 | FOC4326 |
| Peptid markers |  |  |  |  |  |  |  |  |  |  |  |  |  |  |  |  |  |  |  |  |  |
| COL1A1 918-957-RW | □ |  |  |  |  |  |  |  |  |  |  |  |  |  |  |  |  |  |  | □ |  |
| COL1A1 918-957-BW | ○ | ○ | ○ | ○ | ○ | ○ | ○ |  |  | ○ |  |  | ○ | ○ | ○ | ○ |  | ○ | ○ | ○ | ○ |
| COL1A1 1170-1213-RW | □ |  |  |  |  |  |  |  |  |  | □ |  |  |  |  |  |  |  |  | □ |  |
| COL1A1 1170-1213-BW |  | ○ |  | ○ | ○ | ○ | ○ |  | ○ | ○ |  | ○ | ○ | ○ | ○ | ○ | ○ | ○ | ○ |  | ○ |
| COL1A2 262-280-RW | □ |  |  |  |  |  |  | □ |  |  | □ |  |  |  |  |  |  |  |  | □ |  |
| COL1A2 262-280-BW |  | ○ | ○ | ○ | ○ | ○ | ○ |  | ○ | ○ |  | ○ | ○ | ○ | ○ | ○ | ○ | ○ | ○ |  | ○ |
| B. Non-Collagen peptide marker-based species identification of right and bowhead whales using pFind analysis |  |  |  |  |  |  |  |  |  |  |  |  |  |  |  |  |  |  |  |  |  |
| Peptid markers |  |  |  |  |  |  |  |  |  |  |  |  |  |  |  |  |  |  |  |  |  |
| Prothrombin-RW |  |  |  |  |  |  |  |  |  |  |  |  |  |  |  |  |  |  |  | □ |  |
| Prothrombin-BW |  | ○ |  |  |  |  |  |  | ○ | ○ |  |  | ○ |  | ○ |  |  |  | ○ |  |  |
| Identified species | RW | BW | BW | BW | BW | BW | BW | RW | BW | BW | RW | BW | BW | BW | BW | BW | BW | BW | BW | RW | BW |

### 3.5 Historical Balaenidae specimens from the Gulf of St Lawrence

#### 3.5.1 Species assignments and confidence scores

LC–MS/MS analysis successfully identified all previously unidentified balaenid specimens from the Gulf of St Lawrence that were included in this study. Of the 14 previously unidentified historical specimens, 12 were identified as bowhead whale and two as North Atlantic right whale (Table 4). These identifications support previous findings from the Strait of Belle Isle (McLeod et al., 2008), that the majority of Balaenidae in this region in the past were likely to be bowhead whale, not North Atlantic right whale.

**Table 4.** Identification confidence scoring of 14 case study specimens from the Magdalen Islands, previously identified as Balaenidae.

| Specimen | LC–MS/MS identity | Supporting loci | Score | Confidence |
|---|---|---|---|---|
| FOC4050 | Bowhead whale | Prothrombin + COL1A1 1170–1213 + COL1A2 262–280 | 7 | Very high |
| FOC4051 | Bowhead whale | Prothrombin + COL1A1 1170–1213 + COL1A2 262–280 | 7 | Very high |
| FOC4127 | Bowhead whale | Prothrombin + COL1A1 1170–1213 + COL1A2 262–280 | 7 | Very high |
| FOC4195 | North Atlantic Right whale | Prothrombin + COL1A1 1170–1213 + COL1A2 262–280 | 7 | Very high |
| FOC4212 | Bowhead whale | Prothrombin + COL1A1 1170–1213 + COL1A2 262–280 | 7 | Very high |
| FOC4232 | Bowhead whale | Prothrombin + COL1A1 1170–1213 + COL1A2 262–280 | 7 | Very high |
| FOC4238 | North Atlantic Right whale | COL1A1 1170–1213 + COL1A2 262–280 | 5 | High |
| FOC4271 | Bowhead whale | Prothrombin + COL1A1 1170–1213 + COL1A2 262–280 | 7 | Very high |
| FOC4273 | Bowhead whale | Prothrombin + COL1A1 1170–1213 + COL1A2 262–280 | 7 | Very high |
| FOC4274 | Bowhead whale | Prothrombin + COL1A1 1170–1213 + COL1A2 262–280 | 7 | Very high |
| FOC4275 | Bowhead whale | Prothrombin + COL1A1 1170–1213 + COL1A2 262–280 | 7 | Very high |
| FOC4277 | Bowhead whale | Prothrombin + COL1A1 1170–1213 + COL1A2 262–280 | 7 | Very high |
| FOC4282 | Bowhead whale | Prothrombin + COL1A1 1170–1213 + COL1A2 262–280 | 7 | Very high |
| FOC4291 | Bowhead whale | COL1A1 1170–1213 only | 3 | Moderate |

The species assignments were supported primarily by concordant collagen and prothrombin peptide markers, although in two cases (FOC4238, FOC4291) identification depended on collagen markers alone. Bowhead whale specimens consistently (11 out of 12) exhibited the bowhead-associated prothrombin SAP together with the COL1A2 262–280 and COL1A1 1170–1213 bowhead-associated collagen variants. The exception (FOC4291) was identified based on COL1A1 1170–1213 alone. Of the two previously unidentified North Atlantic right whale specimens, one (FOC4195) exhibited the right whale-associated prothrombin SAP together with collagen-derived markers COL1A2 262-280 and COL1A1 1170-1213, whereas the other (FOC4238) showed only the the relevant COL1A2 262–280 and COL1A1 1170–1213 markers.

Application of the weighted multi-locus confidence framework revealed some variation in assignment confidence among the previously unidentified historical specimens depending on the number and specificity of concordant peptide markers recovered (Table 4). The highest-confidence North Atlantic right whale specimen was FOC4195, with a cumulative confidence score of 7 (“very high” confidence). All bowhead whale specimens except FOC4291 also yielded confidence assignment scores of 7. High-confidence assignment (a score of 5) was associated with North Atlantic right whale specimen FOC4238, supported exclusively by two collagen-derived loci. Bowhead FOC4291, being supported exclusively by the COL1A1 1170–1213 marker, accrued a moderate-confidence assignment of 3.

#### 3.5.2 Radiocarbon chronologies and temporal distributions

Radiocarbon dating of the bowhead whale specimens from the Gulf of St Lawrence yielded calibrated age estimates spanning more than four millennia (Figure 1). The earliest specimen (from the Magdalen Islands) dated to ca. 3111–2726 cal BCE, whereas the most recent pre-modern specimens (from the Strait of Belle Isle) dated to the sixteenth to nineteenth centuries cal CE by radiocarbon. Overall, bowhead whale specimens exhibited a broad temporal distribution extending from the late fourth/early third millennium BCE through to the early modern period, with temporally discrete concentrations of dates rather than a uniform distribution through time. Together, these results support a long-term bowhead whale presence in the Gulf of St Lawrence region spanning multiple millennia (Figure 1). Conversely, the two North Atlantic right whales, both from the Magdalen Islands, post-date 1950.

**Figure 1.**
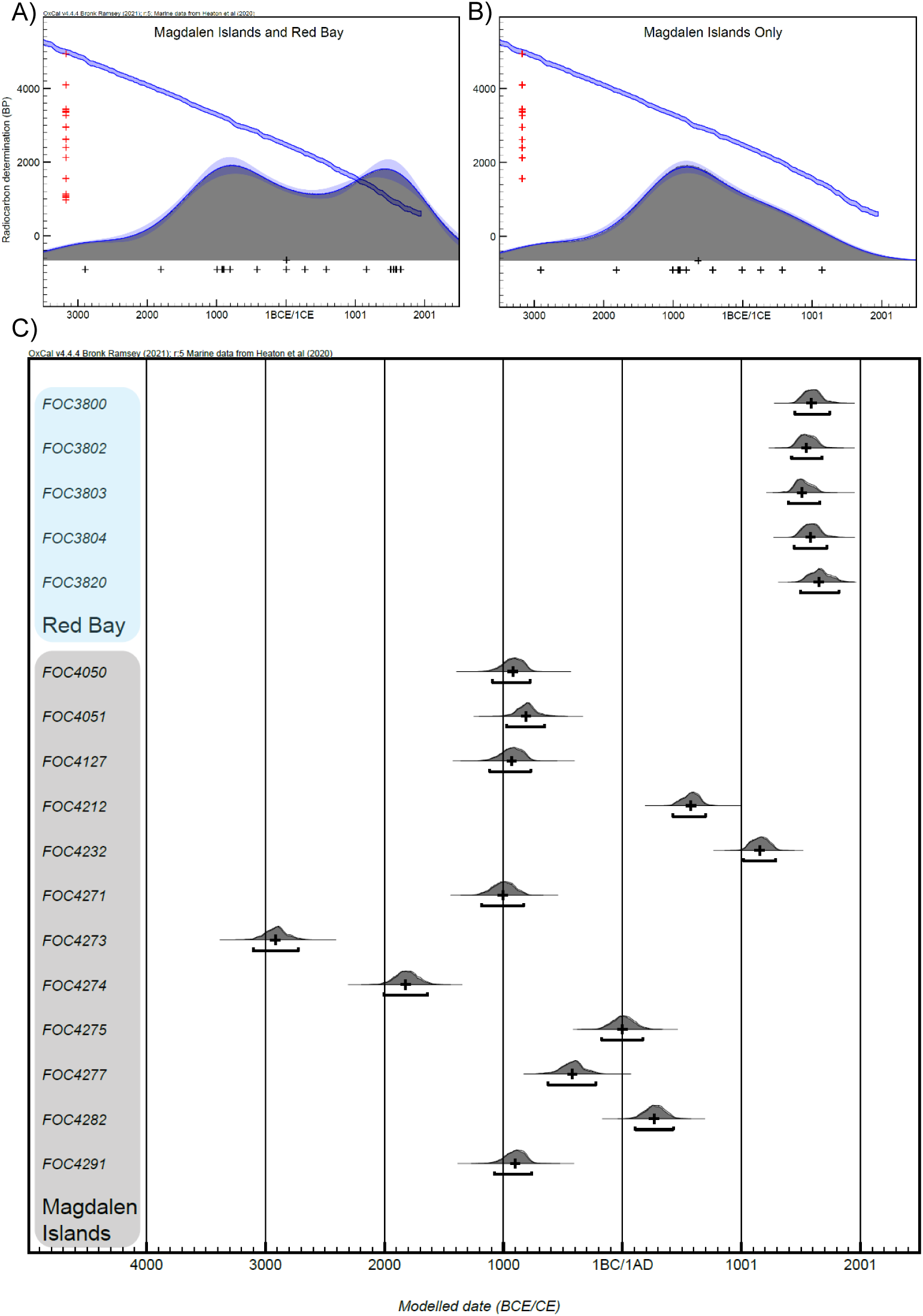
Calibrated radiocarbon dates and OxCal KDE models for bowhead whale specimens from the Gulf of St Lawrence. (A) Kernel density estimation (KDE) model for bowhead whale specimens from the Gulf of St Lawrence, including Magdalen Islands and Strait of Belle Isle/Red Bay samples. Dates were calibrated and modelled in OxCal v4.4 using the Marine20 calibration curve (Heaton et al., 2020) and a regional taxon-specific marine reservoir correction. Cross symbols along the x-axis indicate individual calibrated radiocarbon determinations. (B) KDE model for Magdalen Islands bowhead whale specimens only, generated using the same calibration parameters. (C) Calibrated radiocarbon probability distributions for individual bowhead whale specimens from the Magdalen Islands assemblage and Strait of Belle Isle/Red Bay. Magdalen Islands specimens are shown in the grey-shaded group and Strait of Belle Isle/Red Bay specimens in the blue-shaded group. Probability distributions are shown with 95.4% highest posterior density intervals and reported in cal BCE/CE.

In the Magdalen Islands, calibrated dates for the bowhead whales ranged from 3111–2726 cal BCE to 1014–1284 cal CE, with multiple specimens clustering between approximately ca. 1014 and 818 cal BCE and median estimates centred around ∼900 cal BCE (Figures 1B, 2A). Other bowhead specimens were distributed across the third to second millennia BCE, the first millennium BCE, and the first to second millennia CE, indicating deposition over an extended period at lower rates. Because there was no *a priori* expectation of temporal aggregation within the assemblage, observed and expected date distributions were also compared to evaluate whether deposition was consistent with random temporal sampling through time (Figure 3). The observed distribution showed a stronger concentration of specimens around ∼900 cal BCE than expected under this model, consistent with an aggregated depositional process such as repeated use of a locality, targeted whale exploitation, or a mass-stranding event, although these alternatives cannot be distinguished from the radiocarbon data.

**Figure 2.**
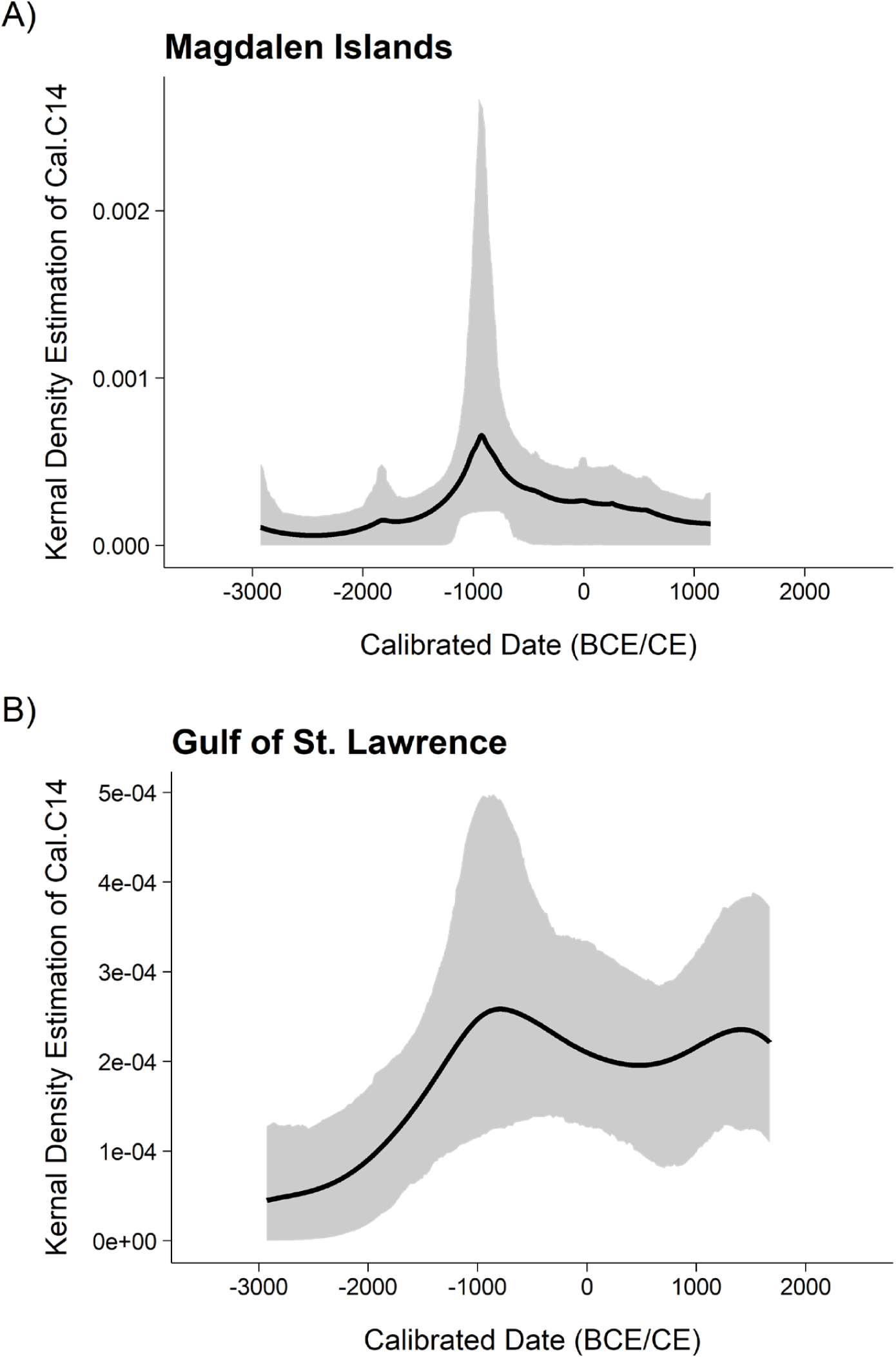
Bootstrapped kernel density estimates of calibrated bowhead whale radiocarbon dates. (A) Kernel density estimate for Magdalen Islands bowhead whale specimens generated in R using 10,000 Monte Carlo simulations. The black line represents the estimated density of calibrated dates, and the grey envelope indicates the bootstrapped 95% confidence interval. (B) Kernel density estimate for the combined Gulf of St Lawrence bowhead whale dataset, including Magdalen Islands and Strait of Belle Isle/Red Bay specimens, generated using the same procedure.

**Figure 3.**
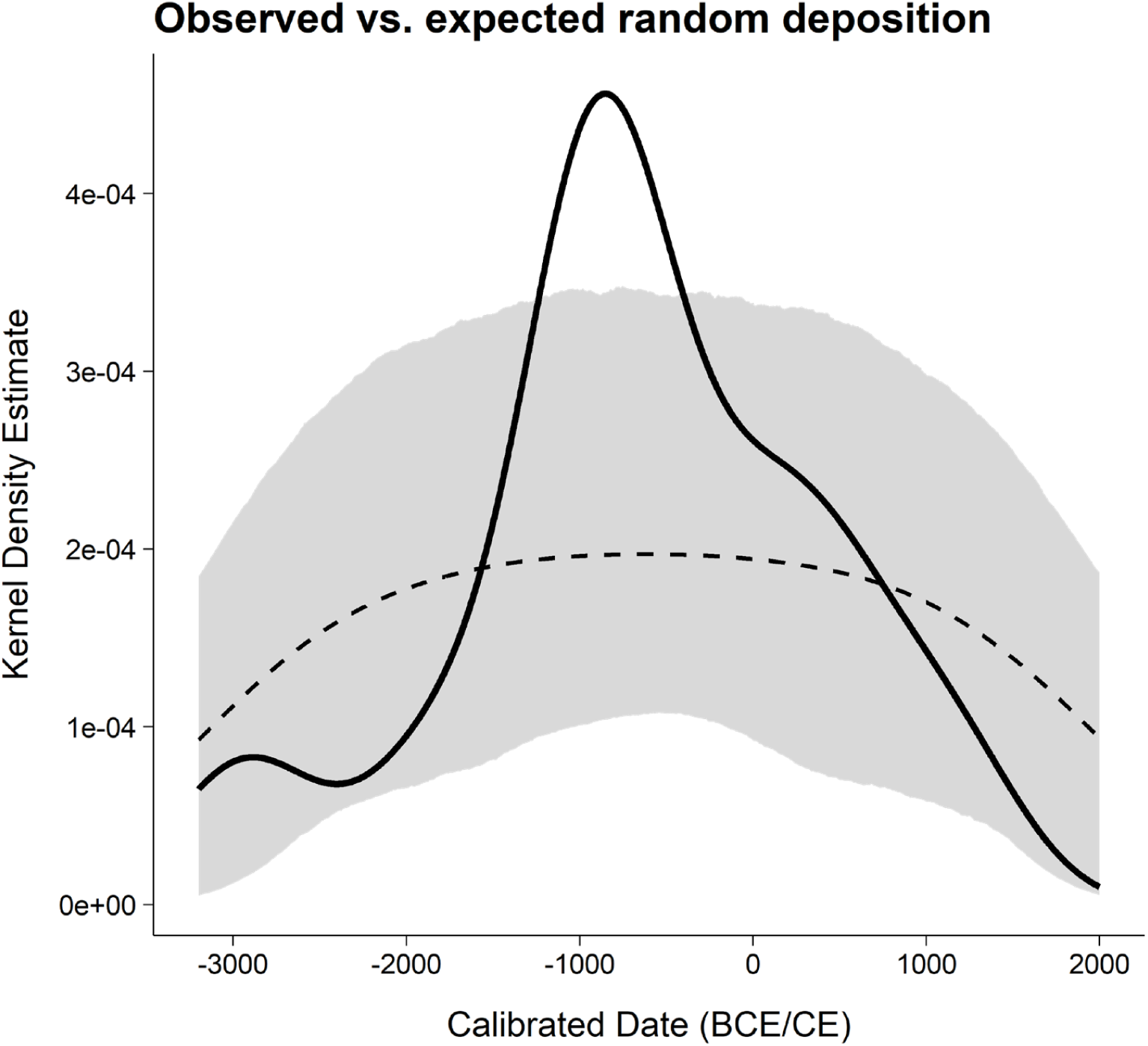
Observed and expected temporal distributions of Magdalen Islands bowhead whale radiocarbon dates. Comparison of the observed temporal distribution of calibrated Magdalen Islands bowhead whale dates with the expected distribution under random temporal sampling. The solid line represents the observed distribution, the dashed line represents the expected distribution, and the grey envelope indicates the simulated confidence interval. This comparison was used to assess whether dated specimens were temporally aggregated relative to a random deposition model.

In contrast, the Red Bay specimens were more tightly clustered, ranging between 1411–1665 and 1507–1840 cal CE, with median estimates between 1526 and 1673 cal CE (Figures 1C, 2B). These dates overlap documented Basque whaling in the Strait of Belle Isle, as expected given both their find context and that they were used to calculate the ΔR value employed for radiocarbon calibration (see above).

KDEs further supported this temporal structure, identifying two principal density concentrations in the combined Gulf of St Lawrence dataset: one centred around ∼900 cal BCE, driven by the Magdalen Islands specimens, and a second around ∼1500 cal CE, driven by the Strait of Belle Isle samples (Figure 1A, 2; note that the modes differ slightly between the OxCal and bootstrapped KDEs, and depending on if the different locations are modelled separately or together). These results indicate at least two temporally distinct periods of elevated bowhead whale deposition within the regional dataset.

## 4 Discussion

This study demonstrates that incorporation of non-collagenous peptide markers into LC–MS/MS workflows facilitates genus or species-level discrimination between the closely related balaenid taxa, the bowhead whale (*Balaena mysticetus*) and the North Atlantic right whale (*Eubalaena glacialis*), which cannot be distinguished using collagen peptide mass fingerprinting alone. Among the evaluated loci, a newly assessed prothrombin-derived peptide containing a glutamic acid/aspartic acid substitution provided robust and reproducible taxonomic discrimination across both modern and historical material, highlighting the potential of non-collagenous proteins. In parallel, evaluation of previously proposed collagen peptide markers demonstrated that some but not all reported loci are consistently diagnostic, with the COL1A1 918–957 marker exhibiting overlapping peptide variants between species and therefore reduced taxonomic reliability in degraded historical material. Finally, although MaxQuant and pFind differed in peptide recovery characteristics, both search engines produced concordant specimen-level species assignments, indicating that the overall taxonomic interpretations are robust to analytical platform choice.

Our findings are particularly important for archaeological and museum specimens, including worked artefacts and highly fragmented subfossil remains, where destructive sampling is often restricted due to specimen rarity, fragility, or cultural significance. The analytical workflow developed here required less than 1 mg of protein extract per specimen, demonstrating that genus or species-level palaeoproteomic identification can be achieved from extremely small quantities of material. This pipeline could be tailored to other historical/archaeological ecofacts or artefacts where taxonomic identifications may only be achievable at low taxonomic resolution because of morphological similarity or fragmentation/manufacturing modification. Distinguishing artefacts of red deer (*Cervus elaphus*) versus moose/elk (*Alces alces*) antler provides one canonical example (Rey-Iglesia et al., 2023; Trolle Jensen et al., 2020; Xue et al., 2022). Genus or species-level identification by LC-MS/MS can be especially important for understanding worked osseous materials, including bone, antler and ivory artefacts (e.g. Dekker et al., 2024; Gilbert et al., 2024; Kumazawa et al., 2026).

### 4.1 Magdalen Islands case study: Historical Balaenidae specimens from the Gulf of St Lawrence

In addition to developing a minimally destructive LC–MS/MS workflow for distinguishing bowhead whale and North Atlantic right whale, this study combined palaeoproteomic and radiocarbon analyses to investigate the long-term historical presence of balaenid whales in the Gulf of St Lawrence, Canada. Accurate discrimination between these closely related taxa is particularly important because both species experienced severe population declines following intensive commercial whaling and are currently classified as endangered or vulnerable in some regions (Kovacs et al., 2020), and critically endangered (Braulik et al., 2023; Vanderlaan et al., 2025), respectively. Moreover, archaeological whale remains are frequently highly fragmented or morphologically ambiguous, limiting genus and species-level identification using conventional zooarchaeological approaches alone (van den Hurk et al., 2023).

The three European control specimens originating from the Netherlands (FOC3489), Spain (FOC3429), and Portugal (FOC3896) were all re-identified as North Atlantic right whale using LC-MS/MS, consistent with historical and zooarchaeological evidence indicating that western European coastal waters formed part of the historical range of this species (Rodrigues et al., 2018), and confirming previous morphological (FOC3489) and genetic (FOC3489, FOC3429) identifications by van den Hurk et al. (2023) and Sigfúsdóttir (2024). In contrast, the relatively warm waters of western Europe would likely have been unsuitable for the more Arctic-adapted bowhead whale. Although Europe was not a focal case study in this instance, we hypothesise that the majority of European Balaenidae specimens from the southern North Sea region and Mediterranean are likely to belong to right whale and not bowhead whale, although long-distance transport or drift of bowhead whale material from further North are plausible (see van den Hurk et al., 2023), and continued molecular confirmation of taxa is critical to our understanding of past species distributions, especially over time scales that entailed major climate change (Foote et al. 2013).

Only four Gulf of St Lawrence specimens in our dataset, two previously unidentified (FOC4195, FOC4238) and two controls (FOC4192, FOC4319), were identified as North Atlantic right whale. All yielded post-1950 dates. Thus there is no evidence for pre-modern occurrence of this species within the analysed material. This pattern is consistent with contemporary species distributions, as North Atlantic right whales are now regularly observed in the Gulf and adjacent waters following a northward shift in migratory range associated with climate-driven habitat change (Durette-Morin et al., 2022). Colder and more extensive sea-ice conditions during parts of the late Holocene (Matzerath et al., 2026; Wu et al., 2022) may have reduced habitat suitability for this species in earlier periods.

All remaining Gulf of St Lawrence specimens, including material from the Magdalen Islands, were identified as bowhead whale. Median radiocarbon dates spanning from 2927 BCE to 1673 CE indicate a prolonged bowhead whale presence within the region (Figure 1; Supplementary Table S1), substantially south of the species’ present-day range (Westbury et al., 2026). This discovery supports previous findings from the Strait of Belle Isle and the northern shore of the Gulf of St Lawrence, where bowhead whales were identified at sixteenth- to seventeenth-century whaling sites (McLeod et al., 2008; Solazzo & Fitzhugh, 2024). KDEs identified two principal concentrations of bowhead whale dates within the combined Gulf of St Lawrence dataset: one centred around ∼900 cal BCE, driven by the Magdalen Islands specimens, and a second around ∼1500 cal CE, driven by the Strait of Belle Isle samples (Figures 1A, 2). The earlier concentration is notable because of its chronology *vis-a-vis* archaeological developments in the region. It falls near the beginning of the Groswater Pre-Dorset archaeological period in Labrador, Newfoundland and the Quebec Lower North Shore (i.e. the northern border of the Gulf of St Lawrence), which was associated with maritime-oriented settlement, fishing, sea-mammal use and coastal mobility (Betts and Hrynick 2021). It is also broadly concurrent with the beginning of the Maritime Woodland archaeological period, known for the exploitation of marine resources elsewhere around the Gulf of St Lawrence (Black 2017). Although the Magdalen Islands specimens cannot be attributed to human utilisation from radiocarbon dates alone, their clustering indicates that bowhead whale deposition was not evenly distributed through time. Instead, the ∼900 cal BCE concentration suggests a discrete episode of elevated deposition, potentially reflecting repeated use of whale-rich shorelines, opportunistic exploitation of stranded whales or more direct engagement with coastal whale resources during a period of cultural and environmental transition. Human occupation of the islands by Indigenous peoples at this time, probably for the exploitation of marine resources, is known from archaeological evidence (McCaffrey 2020). The subsequent dates are more dispersed until the sixteenth–seventeenth century CE concentration associated with Basque whaling in the Strait of Belle Isle. Because radiocarbon data alone cannot distinguish natural stranding, scavenging or active procurement, the Magdalen Islands peak is best interpreted as evidence for increased bowhead whale deposition rather than direct evidence for organised whaling.

In contrast, the later concentration around ∼1500 cal CE reflects both sampling structure and historical whale exploitation, as five bowhead whale specimens derive from the Strait of Belle Isle region, at the Basque whaling station at Red Bay, Labrador. Red Bay was a major centre of sixteenth- and seventeenth-century shore-based whaling, with extensive archaeological evidence for whale processing and large assemblages of bowhead whale remains (McLeod et al., 2008).

Taken together, these findings extend the documented historical range of bowhead whales into the southern Gulf of St Lawrence, where they were present from at least the late fourth/early third millennium BCE. By contrast, the absence of pre-modern North Atlantic right whale specimens suggests that regular occurrence of this species in the Gulf may be comparatively recent, potentially linked to regional oceanographic change and post-industrial warming in the Northwest Atlantic (cf. Durette-Morin et al., 2022). The bowhead whale chronology indicates long-term regional presence through time but also identifies discrete intervals of increased deposition. The assemblage is therefore best interpreted not as a uniform background signal or a single depositional event, but as a multi-phase record combining persistent bowhead whale occurrence with episodic increases in deposition. Although current sample sizes remain limited, additional radiocarbon and palaeoproteomic analyses from the wider Gulf of St Lawrence will allow the spatial and temporal robustness of these patterns to be tested. More broadly, this study demonstrates the value of integrating minimally destructive LC–MS/MS palaeoproteomics with radiocarbon chronologies to reconstruct long-term ecological baselines and human–whale interactions where morphological identification is ambiguous.

### 4.2 Implications for radiocarbon dating and future paleoecological research

The positive bowhead whale ΔR estimated here (+155 ± 35 14C years), derived from approximate-age Red Bay samples dated to AD 1536–1632 (Table S1), is consistent with both bowhead whale ecology and the carbon dynamics of high-latitude, ice-influenced shelf systems (Pearce et al., 2023). Bowhead whales are Arctic and subarctic specialists that feed primarily on zooplankton, often in shallow coastal and sea-ice-influenced habitats (Lowry, 1993). These food webs may incorporate 14C-depleted carbon where restricted air–sea exchange, stratification, advection of older waters, benthic–pelagic coupling, or coastal carbon inputs decouple local production from the global marine reservoir represented by Marine20 (Alves et al., 2018; Pearce et al., 2023). Such processes would increase the apparent radiocarbon age of bowhead whale tissues relative to the global marine calibration curve.

This pattern contrasts with more pelagic rorquals, which typically exploit dense prey fields in offshore or shelf-edge habitats and often at greater depths than bowhead whales (Goldbogen et al., 2012; Guilpin et al., 2019). Differences in habitat use, prey base and water-mass exposure may therefore generate divergent reservoir offsets among baleen whales. Applying the rorqual-derived offset of ca. −188 14C years reported by Buss et al. (under review) rather than the bowhead-specific value of ca. +155 14C years estimated here, would shift the correction by 343 radiocarbon years and alter calibrated age estimates. Thus reservoir corrections cannot be transferred uncritically among baleen whale taxa, which supports the use of species- and region-specific ΔR estimates for marine mammal chronologies.

### 4.3 Limitations and future directions for non-collagenous peptide identification

Although collagen-derived peptides remain central to palaeoproteomic species identification because of their abundance and long-term preservation, our results demonstrate that non-collagenous proteins (NCPs) can provide taxonomic resolution among closely related taxa. In particular, the prothrombin-derived SAP consistently differentiated bowhead whale and North Atlantic right whale across both modern and historical material. These findings highlight the potential for targeted LC–MS/MS workflows to overcome limitations of ZooMS and collagen-only approaches in taxa characterised by highly conserved collagen sequences (Buckley, 2016; Procopio, Hopkins, et al., 2021).

At the same time, this study also illustrates several challenges associated with genus and species-level identification using LC–MS/MS in historical material. Protein recovery is influenced by preservation state, peptide abundance, database completeness, and search-engine behaviour, all of which can affect marker detectability and taxonomic interpretation (Elnaggar et al., 2022). These limitations were particularly evident for several candidate non-collagenous markers, including serotransferrin-like and unconventional myosin-X (see also Solazzo & Fitzhugh, 2024). The low recovery of the serotransferrin-like marker, and the incomplete validation of unconventional myosin-X across available bowhead whale genomic resources, prevented their treatment as primary diagnostic loci. The low recovery of the serotransferrin-like marker illustrates an important challenge in historical palaeoproteomics: the presence of a peptide sequence within a reference genome does not necessarily guarantee recovery within ancient proteomic datasets.

Importantly, our findings also demonstrate that not all previously proposed collagen markers are diagnostic across Balaenidae. The COL1A1 918–957 locus, previously suggested to discriminate between bowhead and right whales (Dekker et al., 2024), was recovered in both taxa in the present study. This observation highlights the benefit of validating peptide markers across larger comparative datasets before applying them to species assignments, particularly in degraded historical material where peptide drop-out may lead to erroneous taxonomic inference if only a single marker is recovered. Biological explanations including retained ancestral polymorphism, lineage-specific variation, or low-frequency hybridisation between balaenid taxa remain possible (Lo et al., 2025) and warrant further genomic investigation. As noted above, proline hydroxylation/oxidation may also be a complicating factor for this peptide.

Differences between MaxQuant and pFind further demonstrate that proteomic search-engine selection can influence peptide detection, particularly for degraded historical proteins. MaxQuant more consistently detected validated non-collagenous markers such as prothrombin. The importance of complementary search approaches for improving peptide-based identifications in complex archaeological and subfossil datasets has previously been highlighted by Elnaggar et al. (2022) and our results further support the value of combining multiple analytical workflows when working with degraded proteins. The differing performance of MaxQuant and pFind likely reflects differences in digestion handling, search-space architecture, and peptide scoring behaviour. Previous benchmarking studies have shown that narrow-search platforms such as MaxQuant and Mascot often perform consistently across a wide range of digestion conditions, whereas open-search systems such as pFind can exhibit increased variability, particularly under semi-tryptic or non-specific search conditions (Freestone et al., 2024). However, in the present study, implementation of semi-specific tryptic searches within MaxQuant improved peptide recovery in degraded historical specimens (FOC4238, FOC4291), indicating that even narrow-search workflows remain sensitive to enzyme specificity settings. Conversely, open-pFind has been shown to recover non-tryptic peptides even during nominally tryptic searches (Chi et al., 2018). Together, these complementary recovery patterns suggest that multi-engine analytical workflows may improve robustness of palaeoproteomic species identification, especially when analysing low-abundance or partially degraded proteins.

More broadly, our results support increasing exploration of NCPs for historical species identification beyond Balaenidae. Candidate proteins including alpha-2-HS-glycoprotein (fetuin-A), biglycan, chondroadherin, pigment epithelium-derived factor (PEDF), and lumican have previously been identified in archaeological bone and may represent promising future taxonomic markers (Ntasi et al., 2022), although their abundance and preservation in cetacean bone remain poorly characterised, they could provide future candidate markers for exploration.

It is worth noting that the present study necessarily relies on relatively small sample numbers, particularly for North Atlantic right whale. It is plausible that multiple loci for each marker may be present within a given species but at comparatively low frequencies. Therefore, as comparative palaeoproteomic datasets expand, additional intra-specific peptide variation may emerge within both collagenous and non-collagenous loci. We provided an example of this with the unexpected occurrence of the COL1A1 918–957 bowhead-associated variant within multiple North Atlantic right whale specimens illustrating the importance of validating candidate markers across broader population-level datasets prior to treating individual loci as universally diagnostic. More generally, these results demonstrate why species identification in degraded material should not rely on single peptide markers alone. Instead, robust taxonomic assignments are best supported through concordance among multiple independently validated loci, particularly where peptide drop-out, differential preservation, or incomplete protein recovery may bias interpretation of individual markers. The weighted multi-locus confidence framework applied here was designed to explicitly accommodate these sources of uncertainty by prioritising reproducible markers with demonstrated taxonomic specificity across multiple control specimens.

Because LC–MS/MS-based palaeoproteomics can be performed on extremely small quantities of material (<1 mg of prepared sample in this study), targeted collagenous and non-collagenous peptide analysis offers a minimally destructive alternative for species identification in fragile, rare, or culturally sensitive specimens where conventional destructive biomolecular approaches may be impractical or prohibited. This is particularly important for worked artefacts, curated museum collections, and highly fragmented subfossil remains that cannot readily be sampled for large-scale genomic analyses, or where preliminary biomolecular screening is required to determine species identity before additional destructive genomic, radiocarbon dating, or isotopic sampling can be justified. Beyond Balaenidae, the multi-locus framework developed here has broader applicability to longstanding archaeological taxonomic problems characterised by poor collagen resolution or morphological ambiguity, including identification of worked antler and other osseous artefacts (e.g. Trolle Jensen et al., 2020; Rey-Iglesia et al., 2023) and differentiation of closely related marine mammal taxa in fragmented coastal assemblages (e.g. Buckley et al., 2014; Hofman et al., 2018). Targeted LC–MS/MS workflows have the potential to bridge the gap between traditional ZooMS screening and more resource-intensive genomic approaches by enabling scalable, high-resolution species identification from minimal starting material. While aDNA approaches can provide high taxonomic resolution, they often require well-preserved material and more intensive laboratory and bioinformatic workflows.

### 4.4 Preventing shifting baselines through LC-MS/MS identification of Archaeofauna

The concept of shifting baselines, originally coined in the context of generational shifts in perspectives around the state of the world’s fisheries (Pauly 1995), describes how each generation perceives the degraded state of ecosystems as the normative or natural condition (McClenachan et al., 2012; Navarro et al., 2025). High-resolution LC–MS/MS palaeoproteomics offers a powerful means of counteracting this phenomenon by enabling genus or species-level identification of highly fragmented archaeological and subfossil faunal remains that would otherwise remain taxonomically ambiguous.

In this study we demonstrate how state-of-the-art biomolecular approaches can reveal the historical presence of species in ecosystems (Beschta & Ripple, 2016; Cooper et al., 2025; Fløjgaard et al., 2022). Our results from the central Gulf of St. Lawrence align with previous genetic evidence from Red Bay and other early Basque whaling stations on the Strait of Belle Isle and the northern shore of the Gulf of St Lawrence, showing that bowhead whale, not North Atlantic right whale, was the dominant balaenid represented in subfossil assemblages for this region (McLeod et al., 2008; Solazzo & Fitzhugh, 2024). This observation confirms that historical balaenid remains in the region did not belong to the right whale, as could be wrongly inferred from present-day occurrences in these waters. By extending the temporal scope of species-occurrence data, biomolecular research (via LC-MS/MS) offers a means to correct potential inaccuracies and reframe historical baselines.

The broader significance of this approach extends beyond Balaenidae. Similar challenges occur across archaeological and palaeontological assemblages where fragmented faunal remains cannot be reliably identified morphologically beyond broad taxonomic categories (see above). More generally, archaeological datasets have repeatedly demonstrated how long-term biomolecular and zooarchaeological evidence can transform understanding of ecosystem change. For example, integrated palaeoecological and archaeological investigations in Chesapeake Bay documented declines of oyster populations by as much as 99% between the sixteenth and nineteenth centuries (Rick & Lockwood, 2013), illustrating how historical baselines inferred from archaeological material can reveal ecological losses far exceeding those recognised from modern observations alone.

As biodiversity loss accelerates, empirical reconstructions of past ecosystems become increasingly important for conservation biology, ecological restoration, and understanding long-term human impacts on environments (Beschta & Ripple, 2016; Cooper et al., 2025; Fløjgaard et al., 2022). In this context, LC–MS/MS palaeoproteomics serves not only as an analytical tool for historical species identification, but also as a means of restoring lost ecological context and preventing contemporary depleted ecosystems from becoming accepted as incomplete historical baselines.

## 5 Conclusion

This study demonstrates that incorporation of non-collagenous peptide markers into LC–MS/MS workflows can achieve genus or species-level discrimination among closely related taxa that are otherwise difficult to distinguish using collagen fingerprinting alone. In particular, a prothrombin-derived peptide containing a validated SAP consistently differentiated bowhead whale and North Atlantic right whale across both modern and historical material, while evaluation of previously proposed collagen loci showed that some but not all collagen-derived markers are diagnostic across Balaenidae.

Our results also highlight the importance of multi-locus approaches for robust palaeoproteomic species identification. Reliance on single peptide markers may produce erroneous taxonomic assignments in degraded archaeological material due to peptide drop-out, incomplete protein recovery, or previously unrecognised intra-specific variation. By integrating concordant evidence from both collagenous and non-collagenous loci, the weighted multi-marker framework applied here increased assignment confidence and reduced dependence on any individual peptide marker. Application of this workflow to historical specimens from the Gulf of St. Lawrence further demonstrated that bowhead whales, rather than North Atlantic right whales, dominated regional balaenid assemblages prior to the twentieth century, refining historical ecological baselines derived from modern species distributions. Because LC–MS/MS requires only tiny quantities of starting material, this approach provides a minimally destructive and scalable alternative for species identification in fragmented, rare, or culturally sensitive material; it offers broad potential for applications in biomolecular archaeology, historical ecology, and conservation palaeobiology.

## Acknowledgments

We thank the following individuals and institutions for kindly providing samples: Musée de la mer, the New Brunswick Museum (Donald McAlpine, Matt Stimson), José Antonio Pis Millán (Centro de Experimentación Pesquera, Dirección General de Pesca Marítima, Gobierno del Principado de Asturias, Gijón, Spain), Carlos Nores (Instituto de Recursos Naturales y Ordenación del Territorio, Universidad de Oviedo, Spain), Youri van den Hurk (University of Bergen) and citizen scientists of the Magdalen Islands (Christine Gouin, Gérald Leblanc, Dany Longuépée, and Emilie Miousse). Youri van den Hurk also kindly commented on an early draft of the manuscript. We acknowledge Rui Venâncio and the Municipality of Peniche for access to the sample FOC3896 (PP.02.509) from the Peniche historical whale bone collection, and Nina Vieira from NOVA FCSH for her scientific support and database organization (CHAM strategic project: FCT reference UID/04666/2025). Animesh Sharma kindly conducted pilot MaxQuant runs. We thank Bastiaan Star, Guðný Ýr Sigfúsdóttir, Lydia Furness and Oliver Kersten (University of Oslo) for sharing the aDNA identifications of samples FOC3429 and FOC3896. All samples were taken with permission and moved across relevant borders in accordance with CITES under institutional registrations CA039, DK014, GB014, NL014 and NO007. Sampling at Red Bay was conducted under permits issued by the Provincial Museum of Newfoundland and Labrador, with additional permission from the Parks Canada Red Bay National Historic Site. This project has received funding from the European Research Council (ERC) under the European Union’s Horizon 2020 research and innovation programme (4-OCEANS, grant agreement no. 951649) and an ERC Proof of Concept grant (PReciSe, grant agreement 101212629). Funded by the European Union. Views and opinions expressed are however those of the author(s) only and do not necessarily reflect those of the European Union or the European Research Council Executive Agency. Neither the European Union nor the granting authority can be held responsible for them. We also thank the many other colleagues who have contributed to this work in terms of logistics and/or advice.

## Author Contributions

Mitra Mohammadi Bazargani and Danielle L. Buss conceived the ideas and designed methodology; Mitra Mohammadi Bazargani conducted proteomics wet lab sample preparation; Lars Hagen collected the proteomics data; Mitra Mohammadi Bazargani analysed the proteomics data; Danielle L. Buss conducted the rarefaction analysis. Danielle L. Buss and James H. Barrett analysed the radiocarbon data; Danielle L. Buss, Jean-Simon Richard and Mohsen Falahati-Anbaran sampled whale bones in the Magdalen Islands; Jean-Simon Richard, Brenna A. Frasier and Cristina Brito contributed samples and information regarding the specimens and their contexts; Mohsen Falahati-Anbaran coordinated the sample handling in Trondheim; Eirik Sollid and Danielle L. Buss conducted protein extraction; Mitra Mohammadi Bazargani and Danielle L. Buss led the writing of the manuscript; James H. Barrett supervised the research and edited and contributed to the manuscript; All authors contributed critically to the drafts and gave final approval for publication.

## References

Alves, E. Q., Macario, K., Ascough, P., & Bronk Ramsey, C. (2018). The worldwide marine radiocarbon reservoir effect: Definitions, mechanisms, and prospects. Reviews of Geophysics, 56(1), 278–305.

Beschta, R. L., & Ripple, W. J. (2016). Riparian vegetation recovery in Yellowstone: The first two decades after wolf reintroduction. Biological Conservation, 198, 93–103.

Betts M. W., & Hrynick M. G. (2021). The Archaeology of the Atlantic Northeast. University of Toronto Press, Toronto.

Black, D.W. (2017) Archaeological sea mammal remains from the Maritime Provinces. Journal of the North Atlantic, Special Volume 10, 70–89.

Borchers, H. (2025) pracma: Practical Numerical Math Functions. R package version 2.4.6. 10.32614/CRAN.package.pracma

Braulik, G. T., Taylor, B. L., Minton, G., Notarbartolo di Sciara, G., Collins, T., Rojas-Bracho, L., Crespo, E. A., Ponnampalam, L. S., Double, M. C., & Reeves, R. R. (2023). Red-list status and extinction risk of the world’s whales, dolphins, and porpoises. Conservation Biology, 37(5), e14090.

Bronk Ramsey, C. (2001) Development of the Radiocarbon Program OxCal. Radiocarbon, 43, 355–363.

Bronk Ramsey, C. (2009) Bayesian analysis of radiocarbon dates. Radiocarbon, 51, 337–360.

Bronk Ramsey, C. (2017) Methods of summarising radiocarbon datasets. Radiocarbon, 59, 1809–1833.

Brown, S., Hebestreit, S., Wang, N., Boivin, N., Douka, K., & Korzow Richter, K. (2020). Zooarchaeology by Mass Spectrometry (ZooMS)-Pretreatment protocols for bone material. protocols.io. 10.17504/protocols.io.bf5djq26

Buckley, M. (2016). Species identification of bovine, Ovine and porcine type 1 collagen; Comparing peptide mass fingerprinting and LC-based proteomics methods. International Journal of Molecular Sciences, 17(4), 445.

Buckley, M., Collins, M., Thomas-Oates, J., & Wilson, J. C. (2009). Species identification by analysis of bone collagen using matrix-assisted laser desorption/ionisation time-of-flight mass spectrometry. Rapid Communications in Mass Spectrometry, 23(23), 3843–3854.

Buckley, M., Fraser, S., Herman, J., Melton, N.D., Mulville, J. & Pálsdóttir, A.H. (2014) Species identification of archaeological marine mammals using collagen fingerprinting. Journal of Archaeological Science, 41, 631–641.

Buss, D. (2021). Foraging ecology and population structuring of baleen whales in the western South Atlantic and eastern South Pacific. PhD thesis. University of Cambridge. 10.17863/CAM.85814

Buss, D. L., O’Connell, T. C., Evans, S., Kneale, C., Osborn, J., Atmore, L. M., Sremba, A. L., Baker, C. S., Dunn, M. J., Goodall-Copestake, W. P., Martin, A., Kitchener, A. C., Stowasser, G., Weir, C. R., & Jackson, J. A. (2025). Historical niche partitioning of Baleen whales in the Western South Atlantic. Marine Ecology Progress Series, 765, 115–136 10.3354/meps14887

Buss, D. L., Richard, J.-S., Mohammadi-Bazargani, M. M., van den Hurk, Y., Gourlay, S., Falahati-Anbaran, M., Frasier, B. A., Sollid, E., & Barrett, J. H. (2026). Bones as biocultural fingerprints: reconstructing potential early historic and indigenous whale encounters in the Gulf of St. Lawrence, Canada. Under Review.

Chi, H., Liu, C., Yang, H., Zeng, W.-F., Wu, L., Zhou, W.-J., Wang, R.-M., Niu, X.-N., Ding, Y.-H., Zhang, Y., Wang, Z.-W., Chen, Z.-L., Sun, R.-X., Liu, T., Tan, G.-M., Dong, M.-Q., Xu, P., Zhang, P.-H., & He, S.-M. (2018). Comprehensive identification of peptides in tandem mass spectra using an efficient open search engine. Nature Biotechnology, 36(11), 1059–1061.

Cooper, R. J., Cabrales, S., Freeman, E., Holroyd, E., Wyatt, J., & Tosney, J. (2025). Eurasian beaver (*Castor fiber*) reintroduction: A nutrient mitigation solution for lowland chalk streams? Wetlands, 45(1), 10. 10.1007/s13157-024-01896-3

Coutu, A. N., Taurozzi, A. J., Mackie, M., Jensen, T. Z. T., Collins, M., & Sealy, J. (2021). Palaeoproteomics confirm earliest domesticated sheep in southern Africa ca. 2000 BP. Scientific Reports, 11. 10.1038/s41598-021-85756-8

Dekker, J. A. A., Mylopotamitaki, D., Verbaas, A., Sinet-Mathiot, V., Presslee, S., McCarthy, M. L., Olsen, M. T., Olsen, J. V., van den Hurk, Y., Brattinga, J., & Welker, F. (2024). Palaeoproteomic identification of a whale bone tool from Bronze Age Heiloo, the Netherlands. Peer Community Journal, 4(e81). 10.24072/pcjournal.451

Dorai-Raj, S. & Cascone, A.D. (2026) binom: Binomial Confidence Intervals for Several Parameterizations. R package version 1.1-2. 10.32614/CRAN.package.binom

Durette-Morin, D., Evers, C., Johnson, H. D., Kowarski, K., Delarue, J., Moors-Murphy, H., Maxner, E., Lawson, J. W., & Davies, K. T. A. (2022). The distribution of North Atlantic right whales in Canadian waters from 2015-2017 revealed by passive acoustic monitoring. Frontiers in Marine Science, 9. 10.3389/fmars.2022.976044

Elnaggar, A., Osama, A., Anwar, A. M., Ezzeldin, S., Abou Elhassan, S., Ebeid, H., Leona, M., & Magdeldin, S. (2022). Paleoproteomic profiling for identification of animal skin species in ancient Egyptian archaeological leather using liquid chromatography coupled with tandem mass spectrometry (LC-MS/MS). Heritage Science, 10(1). 10.1186/s40494-022-00816-0

Engels, I., Burnett, A., Robert, P., Pironneau, C., Abrams, G., Bouwmeester, R., Van der Plaetsen, P., Di Modica, K., Otte, M., Straus, L. G., Fischer, V., Bray, F., Mesuere, B., De Groote, I., Deforce, D., Daled, S., & Dhaenens, M. (2025). Classification of collagens via peptide ambiguation, in a paleoproteomic LC-MS/MS-based taxonomic pipeline. Journal of Proteome Research, 24(4), 1907–1925.

Fahrner, M., Kook, L., Fröhlich, K., Biniossek, M. L., & Schilling, O. (2021). A systematic evaluation of semispecific peptide search parameter enables identification of previously Undescribed N-terminal peptides and conserved proteolytic processing in cancer cell lines. Proteomes, 9(2), 26.

Fløjgaard, C., Pedersen, P. B. M., Sandom, C. J., Svenning, J.-C., & Ejrnæs, R. (2022). Exploring a natural baseline for large-herbivore biomass in ecological restoration. Journal of Applied Ecology, 59(1), 18–24.

Foote, A.D., Kaschner, K., Schultze, S.E., Garilao, C., Ho, S.Y., Post, K., … Gilbert, M. T. P. (2013). Ancient DNA reveals that bowhead whale lineages survived Late Pleistocene climate change and habitat shifts. Nature Communications, 4, 1677.

Freestone, J., Noble, W. S., & Keich, U. (2024). Reinvestigating the correctness of decoy-based false discovery rate control in proteomics tandem mass spectrometry. Journal of Proteome Research, 23(6), 1907–1914.

Gaeta, R. (2021). Ancient DNA and paleogenetics: risks and potentiality. Pathologica, 113(2), 141–146.

Gilbert, C., Krupicka, V., Galluzzi, F., Popowich, A., Bathany, K., Claverol, S., Arslanoglu, J., & Tokarski, C. (2024). Species identification of ivory and bone museum objects using minimally invasive proteomics. Science Advances, 10(4), eadi9028.

Giuffrida, M. G., Mazzoli, R., & Pessione, E. (2018). Back to the past: deciphering cultural heritage secrets by protein identification. Applied Microbiology and Biotechnology, 102(13), 5445–5455.

Goldbogen, J. A., Calambokidis, J., Croll, D. A., McKenna, M. F., Oleson, E., Potvin, J., Pyenson, N. D., Schorr, G., Shadwick, R. E., & Tershy, B. R. (2012). Scaling of lunge-feeding performance in rorqual whales: mass-specific energy expenditure increases with body size and progressively limits diving capacity. Functional Ecology, 26(1), 216–226.

Guilpin, M., Lesage, V., McQuinn, I., Goldbogen, J. A., Potvin, J., Jeanniard-du-Dot, T., Doniol-Valcroze, T., Michaud, R., Moisan, M., & Winkler, G. (2019). Foraging energetics and prey density requirements of western North Atlantic blue whales in the Estuary and Gulf of St. Lawrence, Canada. Marine Ecology Progress Series, 625, 205–223.

Heaton, T., Köhler, P., Butzin, M., Bard, E., Reimer, R., Austin, W., Bronk Ramsey, C., Grootes, P., Hughen, K.A., Kromer, B., Reimer, P.J., Adkins, J., Burke, A., Cook, M.S., Olsen, J. & Skinner, L. (2020) Marine20—The marine radiocarbon age calibration curve (0–55,000 cal BP). Radiocarbon, 62, 779–820.

Hofman, C.A., Rick, T.C., Erlandson, J.M., Reeder-Myers, L., Welch, A.J. & Buckley, M. (2018) Collagen fingerprinting and the earliest marine mammal hunting in North America. Scientific Reports, 8, 10014.

Janzen, A., Richter, K. K., Mwebi, O., Brown, S., Onduso, V., Gatwiri, F., Ndiema, E., Katongo, M., Goldstein, S. T., Douka, K., & Boivin, N. (2021). Distinguishing African bovids using Zooarchaeology by Mass Spectrometry (ZooMS): New peptide markers and insights into Iron Age economies in Zambia. PLOS ONE, 16(5), e0251061.

Keane, M., Semeiks, J., Webb, A. E., … de Magalhães, J. P. (2015). Insights into the evolution of longevity from the bowhead whale genome. Cell Reports, 10, 112–120.

Kovacs, K. M., Lydersen, C., Vacquiè-Garcia, J., Shpak, O., Glazov, D., & Heide-Jørgensen, M. P. (2020). The endangered Spitsbergen bowhead whales’ secrets revealed after hundreds of years in hiding. Biology Letters, 16(6), 20200148.

Kumazawa, Y., Mizuno, K., & Taga, Y. (2026). Species identification of elephant and mammoth ivory products by LC-MS analysis of type I collagen marker peptides with a microsampling technique. Talanta, 306(129763), 129763.

Lanigan, L. T., Mackie, M., Feine, S., Hublin, J.-J., Schmitz, R. W., Wilcke, A., Collins, M. J., Cappellini, E., Olsen, J. V., Taurozzi, A. J., & Welker, F. (2020). Multi-protease analysis of Pleistocene bone proteomes. Journal of Proteomics, 228(103889), 103889.

Lo, B.-W., Martinez Real, F., Magg, A., Wise, J. P., Sr, Mundlos, S., & Franchini, P. (2025). Genome-wide demographic analyses of balaenid whales revealed complex history of gene flow associated with past climate oscillation. Genome Biology and Evolution, 17(5), evaf081.

Lowry, L. F. (1993). Foods and feeding ecology. J.J. Burns & J. Montaque (Eds.), The Bowhead Whale (pp. 201–238). Society for Marine Mammalogy.

Magnadóttir, B., Uysal-Onganer, P., Kraev, I., Svansson, V., Hayes, P., & Lange, S. (2020). Deiminated proteins and extracellular vesicles - Novel serum biomarkers in whales and orca. Comparative Biochemistry and Physiology. Part D, Genomics & Proteomics, 34(100676), 100676.

Matzerath, P., Kolling, H. M., Kienast, M., Fahl, K., Stein, R., Gross, F., & Schneider, R. R. (2026). Holocene sea-ice and productivity changes on the Scotian Shelf, NW Atlantic. Quaternary Science Reviews, 372(109694), 109694.

McCaffrey, M. (2020) Atlantic walrus in the Gulf of St. Lawrence: A history of human predation. I. Krupnik & A. L. Crowell (Eds.), Arctic Crashes: People and Animals in the Changing North (pp. 375–399). Smithsonian Scholarly Press.

McClenachan, L., Ferretti, F., & Baum, J. K. (2012). From archives to conservation: why historical data are needed to set baselines for marine animals and ecosystems. Conservation Letters, 5(5), 349–359.

McLeod, B. A., Brown, M. W., Moore, M. J., Stevens, W., Barkham, S. H., Barkham, M., & White, B. N. (2008). Bowhead Whales, and not Right Whales, were the primary target of 16th-to 17th-century Basque whalers in the western North Atlantic. Arctic, 61(1), 61–75.

Navarro, L.M., Armstrong, C.G., Changeux, T., Frisch, D., Gil-Romera, G., Kaim, D., et al. (2025) Integrating historical sources for long-term ecological knowledge and biodiversity conservation. Nature Reviews Biodiversity, 1, 657–670.

Ntasi, G., Palomo, I. R., Marino, G., Piaz, F. D., Cappellini, E., Birolo, L., & Petrone, P. (2022). Molecular signatures written in bone proteins of 79 AD victims from Herculaneum and Pompeii. Scientific Reports, 12(1), 8401.

Oksanen J, S.G., Blanchet F, Kindt R, Legendre P, Minchin P, O’Hara R, Solymos P, Stevens M, Szoecs E, Wagner H, Bedward M, Bolker B, Borcard D, Carvalho G, De Caceres M, Durand S, Evangelista H, Hannigan G, Hill M, Lahti L, Martino C, Ouellette M, Ribeiro Cunha E, Smith T, Stier A, Ter Braak C, Weedon J (2026) vegan: Community Ecology Package. R package version 2.7-6. 10.32614/CRAN.package.vegan

Pääbo, S., Poinar, H., Serre, D., Jaenicke-Despres, V., Hebler, J., Rohland, N., Kuch, M., Krause, J., Vigilant, L., & Hofreiter, M. (2004). Genetic analyses from ancient DNA. Annual Review of Genetics, 38(1), 645–679.

Pearce, C., Özdemir, K. S., Forchhammer, R. C., Detlef, H., & Olsen, J. (2023). The marine reservoir age of Greenland coastal waters. Geochronology, 5, 451–465.

Perez-Riverol, Y., Bai, J., Bandla, C., García-Seisdedos, D., Hewapathirana, S., Kamatchinathan, S., Kundu, D. J., Prakash, A., Frericks-Zipper, A., Eisenacher, M., Walzer, M., Wang, S., Brazma, A., & Vizcaíno, J. A. (2022). The PRIDE database resources in 2022: a hub for mass spectrometry-based proteomics evidences. Nucleic Acids Research, 50(D1), D543–D552.

Procopio, N., Hopkins, R. J. A., Harvey, V. L., & Buckley, M. (2021). Proteome variation with collagen yield in ancient bone. Journal of Proteome Research, 20(3), 1754–1769.

Procopio, N., Mein, C. A., Starace, S., Bonicelli, A., & Williams, A. (2021). Bone diagenesis in short timescales: Insights from an exploratory proteomic analysis. Biology, 10(6), 460.

Rey-Iglesia, A., de Jager, D., Presslee, S., Qvistgaard, S. S., Sindbæk, S. M., & Lorenzen, E. D. (2023). Antlers far and wide: Biomolecular identification of Scandinavian hair combs from Ribe, Denmark, 720–900 CE. Journal of Archaeological Science, 153(105773), 105773.

Richter, K. K., Codlin, M. C., Seabrook, M., & Warinner, C. (2022). A primer for ZooMS applications in archaeology. Proceedings of the National Academy of Sciences, 119(20), e2109323119.

Rick, T. C., & Lockwood, R. (2013). Integrating paleobiology, archeology, and history to inform biological conservation: Paleobiology, archeology, and history. Conservation Biology: The Journal of the Society for Conservation Biology, 27(1), 45–54.

Rodrigues, A. S. L., Charpentier, A., Bernal-Casasola, D., Gardeisen, A., Nores, C., Pis Millán, J. A., McGrath, K., & Speller, C. F. (2018). Forgotten Mediterranean calving grounds of grey and North Atlantic right whales: evidence from Roman archaeological records. Proceedings of the Royal Society B: Biological Sciences 285(1882). 10.1098/rspb.2018.0961

Sawafuji, R., Cappellini, E., Nagaoka, T., Fotakis, A. K., Jersie-Christensen, R. R., Olsen, J. V., Hirata, K., & Ueda, S. (2017). Proteomic profiling of archaeological human bone. Royal Society Open Science, 4(6), 161004.

Shapiro, B., & Hofreiter, M. (2014). A paleogenomic perspective on evolution and gene function: new insights from ancient DNA. Science, 343(6169), 1236573.

Sigfúsdóttir, G.Ý. (2024) Whale Tales: Genetic impacts of human exploitation on the North Atlantic right whale (Eubalaena glacialis) investigated using ancient DNA. Master’s thesis, University of Oslo, Oslo, Norway. https://hdl.handle.net/10852/113407

Solazzo, C., & Fitzhugh, W. W. (2024). Proteomics identify bowhead whale muscle tissue and baleen in cinder residues at a 16th c. Basque whaling site. Journal of Archaeological Science: Reports, 58(104691), 104691.

Steen, H., & Mann, M. (2004). The ABC’s (and XYZ’s) of peptide sequencing. Nature Reviews. Molecular Cell Biology, 5(9), 699–711.

Thermo Fisher Scientific (2022) Pierce™ Trypsin Protease, MS Grade User Guide (Pub. No. MAN0011821, Rev. B.0). Thermo Fisher Scientific, Rockford, IL, USA.

Trolle Jensen, T. Z., Mackie, M., Taurozzi, A. J., Lanigan, L. T., Gundelach, C., Olsen, J., Sørensen, S. A., Collins, M. J., Sørensen, M., & Schroeder, H. (2020). The biomolecular characterization of a finger ring contextually dated to the emergence of the Early Neolithic from Syltholm, Denmark. Royal Society Open Science, 7(1), 191172.

Tyanova, S., Temu, T. & Cox, J. (2016) The MaxQuant computational platform for mass spectrometry-based shotgun proteomics. Nature protocols, 11, 2301–2319.

van den Hurk, Y., Sikström, F., Amkreutz, L., Bleasdale, M., Borvon, A., Ephrem, B., Fernández-Rodríguez, C., Gibbs, H. M. B., Jonsson, L., Lehouck, A., Cedeira, J. M., Meng, S., Monge, R., Moreno, M., Nabais, M., Nores, C., Pis-Millán, J. A., Riddler, I., Schmölcke, U., … Barrett, J. H. (2023). The prelude to industrial whaling: identifying the targets of ancient European whaling using zooarchaeology and collagen mass-peptide fingerprinting. Royal Society Open Science, 10(9), 230741.

Vanderlaan, A. S., Lang, S. L., Sanchez, M., Murphy, M. J., Pisano, O. M., & Christie, K. (2025). Threat assessment for the critically endangered North Atlantic right whale (*Eubalaena glacialis*). https://open-science.canada.ca/server/api/core/bitstreams/e2689b0a-aad2-4fec-a196-2c09abfd1ed8/content

Wadsworth, C., & Buckley, M. (2014). Proteome degradation in fossils: investigating the longevity of protein survival in ancient bone. Rapid Communications in Mass Spectrometry, 28(6), 605–615.

Wadsworth, C., & Buckley, M. (2018). Characterization of proteomes extracted through collagen-based stable isotope and radiocarbon dating methods. Journal of Proteome Research, 17(1), 429–439.

Warinner, C., Korzow Richter, K., & Collins, M. J. (2022). Paleoproteomics. Chemical Reviews, 122(16), 13401–13446.

Welker, F., Soressi, M., Rendu, W., Hublin, J.-J., & Collins, M. (2015). Using ZooMS to identify fragmentary bone from the Late Middle/Early Upper Palaeolithic sequence of Les Cottés, France. Journal of Archaeological Science, 54, 279–286.

Westbury, M. V., Freymueller, N. A., Cabrera, A. A., Bachmann, L., Ferguson, S. H., Heide-Jørgensen, M. P., Kovacs, K. M., Lydersen, C., Shpak, O., Wiig, Ø., Fordham, D. A., & Lorenzen, E. D. (2026). Tracing the drivers of range-wide bowhead whale genomic structure and diversity. Molecular Ecology, 35(10), e70378.

Wu, X., de Vernal, A., Fréchette, B., Moros, M., & Perner, K. (2022). The signal of climate changes over the last two millennia in the Gulf of St. Lawrence, eastern Canada. Quaternary Research, 106, 28–43.

Xue, F., Wang, B., Guo, D.-X., Jiao, Y., Yin, X., Cui, W.-L., Zhou, Q.-Q., Yu, F.-R., & Lin, Y.-Q. (2022). Peptide biomarkers discovery for seven species of deer antler using LC-MS/MS and label-free approach. Molecules, 27(15), 4756.

