## Supplementary Tables and Figure for "Non-collagenous and collagenous peptide markers for distinguishing between closely-related species using LC-MS/MS: A case study in baleen whales"

**Supplementary Table S1.** The samples studied and related metadata, including the origins of the specimens, their roles in the study, taxonomic identifications and radiocarbon dates. North Atlantic right whale is abbreviated NARW.

| Sample | Type of sample | Prior taxonomic identification | Prior identification method | Source of previous identification | Final LC-MS/MS taxonomic identity | Radiocarbon lab code (TRa-) | Radiocarbon date (uncalibrated) | PMC values | $\delta^{13}C$ from AMS | Source of radiocarbon date | Radiocarbon date BCE/CE (Calibrated at 95.4% using the Marine20 curve with delta R) | Median radiocarbon date or historical date BCE/CE (Calibrated using the Marine20 curve with delta R) | Delta R employed | Sample reference | Reference for Delta R | Location | Sample provider | Original specimen identifier | Oslo aDNA number | Frasier Lab aDNA number |
|---|---|---|---|---|---|---|---|---|---|---|---|---|---|---|---|---|---|---|---|---|
| FOC3489 | Historical control | Bowhead | Morphology | Van den Hurk et al. 2023 | Bowhead | 17850 | 585 ± 15 BP | 93.00 ± 0.16 | -15.1 ± 0.1 ‰ | This study | 1833 - 1953 | 1913 | 155 ± 35* | Van den Hurk et al. 2023 | This study | Zeeland, Netherlands | Koninklijk Zeeuwsch Genootschap der Wetenschappen/ Youri van den Hurk | 1316; DAG442 | | |
| FOC3800 | Historical control | Bowhead | DNA | McLeod et al. 2008 | Bowhead | 21501 | 1040 ± 20 BP | 87.87 ± 0.18 | -13.7 ± 2.1 ‰ | This study | 1464 - 1769 | 1606 | 155 ± 35** | McLeod et al. 2008 | This study | Red Bay, Strait of Belle Isle, Canada | Brenna Frasier/Parks Canada | BMMS 34 | | WHB0114 |
| FOC3802 | Historical control | Bowhead | DNA | McLeod et al. 2008 | Bowhead | 21500 | 1095 ± 25 BP | 87.24 ± 0.25 | -16.0 ± 0.2 ‰ | This study | 1432 - 1693 | 1561 | 155 ± 35** | McLeod et al. 2008 | This study | Red Bay, Strait of Belle Isle, Canada | Brenna Frasier/Parks Canada | BMMS 36 | | WHB0116 |
| FOC3803 | Historical control | Bowhead | DNA | McLeod et al. 2008 | Bowhead | 21499 | 1135 ± 20 BP | 86.84 ± 0.20 | -15.5 ± 0.4 ‰ | This study | 1411 - 1665 | 1526 | 155 ± 35** | McLeod et al. 2008 | This study | Red Bay, Strait of Belle Isle, Canada | Brenna Frasier/Parks Canada | BMMS 38 | | WHB0118 |
| FOC3804 | Historical control | Bowhead | DNA | McLeod et al. 2008 | Bowhead | 21498 | 1050 ± 20 BP | 87.76 ± 0.21 | -14.0 ± 1.9 ‰ | This study | 1460 - 1751 | 1598 | 155 ± 35** | McLeod et al. 2008 | This study | Red Bay, Strait of Belle Isle, Canada | Brenna Frasier/Parks Canada | BMMS 41 | | WHB0120 |
| FOC3820 | Historical control | Bowhead | DNA | McLeod et al. 2008 | Bowhead | 21494 | 975 ± 20 BP | 88.58 ± 0.19 | -14.6 ± 1.7 ‰ | This study | 1507 - 1840 | 1673 | 155 ± 35** | McLeod et al. 2008 | This study | Red Bay, Strait of Belle Isle, Canada | Brenna Frasier/Parks Canada | 24M-2004-066 | | WHB0379 |
| FOC4050 | Case study | Unidentified Balaenidae | ZooMS | Buss et al. under review | Bowhead | 21750 | 3373 ± 22 BP | 65.71 ± 0.18 | -15.2 ± 1.1 ‰ | Buss et al. under review | -1107 - -782 | -930 | 155 ± 35 | Buss et al. under review | This study | Magdalen Islands, Gulf of St. Lawrence, Québec, Canada | Jean-Simon Richard | NA | | |
| FOC4051 | Case study | Unidentified Balaenidae | ZooMS | Buss et al. under review | Bowhead | 22133 | 3269 ± 18 BP | 66.57 ± 0.14 | -13.6 ± 0.3 ‰ | Buss et al. under review | -977 - -663 | -818 | 155 ± 35 | Buss et al. under review | This study | Magdalen Islands, Gulf of St. Lawrence, Québec, Canada | Emilie Miousse | NA | | |
| FOC4127 | Case study | Unidentified Balaenidae | ZooMS | Buss et al. under review | Bowhead | 22134 | 3380 ± 35 BP | 65.65 ± 0.27 | -14.1 ± 1.7 ‰ | Buss et al. under review | -1126 - -776 | -939 | 155 ± 35 | Buss et al. under review | This study | Magdalen Islands, Gulf of St. Lawrence, Québec, Canada | Christine Gouin | NA | | |
| FOC4212 | Case study | Unidentified Balaenidae | ZooMS | Buss et al. under review | Bowhead | 22165 | 2125 ± 15 BP | 76.77 ± 0.13 | -14.0 ± 0.4 ‰ | Buss et al. under review | 421 - 696 | 569 | 155 ± 35 | Buss et al. under review | This study | Magdalen Islands, Gulf of St. Lawrence, Québec, Canada | Jean-Simon Richard | NA | | |
| FOC4232 | Case study | Unidentified Balaenidae | ZooMS | Buss et al. under review | Bowhead | 22142 | 1556 ± 17 BP | 82.39 ± 0.17 | -13.9 ± 0.8 ‰ | Buss et al. under review | 1014 - 1284 | 1147 | 155 ± 35 | Buss et al. under review | This study | Magdalen Islands, Gulf of St. Lawrence, Québec, Canada | Jean-Simon Richard | NA | | |
| FOC4271 | Case study | Unidentified Balaenidae | ZooMS | Buss et al. under review | Bowhead | 21753 | 3440 ± 17 BP | 65.16 ± 0.13 | -14.7 ± 0.2 ‰ | Buss et al. under review | -1194 - -837 | -1014 | 155 ± 35 | Buss et al. under review | This study | Magdalen Islands, Gulf of St. Lawrence, Québec, Canada | Jean-Simon Richard | NA | | |
| FOC4273 | Case study | Unidentified Balaenidae | ZooMS | Buss et al. under review | Bowhead | 21755 | 4940 ± 20 BP | 54.06 ± 0.12 | -14.8 ± 0.9 ‰ | Buss et al. under review | -3111 - -2726 | -2927 | 155 ± 35 | Buss et al. under review | This study | Magdalen Islands, Gulf of St. Lawrence, Québec, Canada | Jean-Simon Richard | NA | | |
| FOC4274 | Case study | Unidentified Balaenidae | ZooMS | Buss et al. under review | Bowhead | 21756 | 4100 ± 20 BP | 60.01 ± 0.14 | -13.7 ± 4.1 ‰ | Buss et al. under review | -2017 - -1647 | -1831 | 155 ± 35 | Buss et al. under review | This study | Magdalen Islands, Gulf of St. Lawrence, Québec, Canada | Jean-Simon Richard | NA | | |
| FOC4275 | Case study | Unidentified Balaenidae | ZooMS | Buss et al. under review | Bowhead | 22167 | 2618 ± 20 BP | 72.19 ± 0.17 | -13.9 ± 0.4 ‰ | Buss et al. under review | -184 - 165 | -9 | 155 ± 35 | Buss et al. under review | This study | Magdalen Islands, Gulf of St. Lawrence, Québec, Canada | Jean-Simon Richard | NA | | |
| FOC4277 | Case study | Unidentified Balaenidae | ZooMS | Buss et al. under review | Bowhead | 22169 | 2952 ± 22 BP | 69.25 ± 0.18 | -13.6 ± 0.6 ‰ | Buss et al. under review | -654 - -238 | -432 | 155 ± 35 | Buss et al. under review | This study | Magdalen Islands, Gulf of St. Lawrence, Québec, Canada | Dany Longuépée | NA | | |
| FOC4282 | Case study | Unidentified Balaenidae | ZooMS | Buss et al. under review | Bowhead | 22174 | 2399 ± 15 BP | 74.18 ± 0.12 | -14.6 ± 0.2 ‰ | Buss et al. under review | -97 - 420 | 258 | 155 ± 35 | Buss et al. under review | This study | Magdalen Islands, Gulf of St. Lawrence, Québec, Canada | Dany Longuépée | NA | | |
| FOC4291 | Case study | Unidentified Balaenidae | ZooMS | Buss et al. under review | Bowhead | 21763 | 3357 ± 22 BP | 65.84 ± 0.17 | -13.4 ± 0.3 ‰ | Buss et al. under review | -1094 - -768 | -910 | 155 ± 35 | Buss et al. under review | This study | Magdalen Islands, Gulf of St. Lawrence, Québec, Canada | Jean-Simon Richard | NA | | |
| FOC4326 | Modern control | Bowhead | Modern stranding | New Brunswick Museum, Canada; Ledwell et al. 2007 | Bowhead | NA | NA | NA | NA | NA | NA | 2005 | NA | This study | NA | Witless Bay Point on the southern shore of the Avalon Penninsula, Newfoundland | New Brunswick Museum | NBM-MA-011485 | | |
| FOC3429 | Historical control | NARW | DNA | Sigfúsdóttir 2024 | NARW | 23069 | 775 ± 20 BP | 90.81 ± 0.22 | -15.1 ± 1.3 ‰ | This study | 1375 - 1795 | 1559 | -158 ± 88 | Van den Hurk et al. 2023 | Buss et al. under review | Port of Candás, Llanes, Asturias, Spain (43.590 -5.764) | University of Oviedo / José Antonio Pis Millán / Carlos Nores / Youri van den Hurk | 2021-05-17 (004); DAG276 | WHL035 | |
| FOC3896 | Historical control | NARW | DNA | Sigfúsdóttir 2024 | NARW | 21491 | 1185 ± 15 BP | 86.31 ± 0.14 | -17.1 ± 0.5 ‰ | This study | 1019 - 1414 | 1223 | -158 ± 88 | This study | Buss et al. under review | Porto de Pesca, Peniche, Portugal | Municipality of Peniche / Rui Venâncio / Nina Vieira / Cristina Brito | PP.02.509 | WHL022 | |
| FOC4192 | Historical control | NARW | Morphology | Buss et al. under review | NARW | 22137 | > 1950 AD | 102.95 ± 0.18 | -15.8 ± 1.1 ‰ | Buss et al. under review | Post-1950 | Post-1950 | NA | Buss et al. under review | NA | Magdalen Islands, Gulf of St. Lawrence, Québec, Canada | Musée de la mer | NA | | |
| FOC4195 | Case study | Unidentified Balaenidae | ZooMS | Buss et al. under review | NARW | 22176 | > 1950 AD | 102.55 ± 0.14 | -15.2 ± 0.5 ‰ | Buss et al. under review | Post-1950 | Post-1950 | NA | Buss et al. under review | NA | Magdalen Islands, Gulf of St. Lawrence, Québec, Canada | Gerald Leblanc | NA | | |
| FOC4238 | Case study | Unidentified Balaenidae | ZooMS | Buss et al. under review | NARW | 22148 | > 1950 AD | 102.11 ± 0.15 | -16.4 ± 0.2 ‰ | Buss et al. under review | Post-1950 | Post-1950 | NA | Buss et al. under review | NA | Magdalen Islands, Gulf of St. Lawrence, Québec, Canada | Jean-Simon Richard | NA | | |
| FOC4314 | Modern control | NARW | Modern stranding | New Brunswick Museum, Canada | NARW | NA | NA | NA | NA | NA | NA | 2006 | NA | This study | NA | Roseway Basin, 5 miles off Yarmouth, Nova Scotia, Canada | New Brunswick Museum | NBM-MA-011766 | | |
| FOC4319 | Modern control | NARW | Modern ship strike | New Brunswick Museum, Canada | NARW | NA | NA | NA | NA | NA | NA | 2019 | NA | This study | NA | Gulf of St. Lawrence, New Brunswick (48.25 -63.138) | New Brunswick Museum | NBM-MA-018224 | | |
| FOC4323 | Modern control | NARW | Modern stranding | New Brunswick Museum, Canada | NARW | NA | NA | NA | NA | NA | NA | 1994 | NA | This study | NA | Southern Cove, Kent Island, New Brunswick | New Brunswick Museum | NBM-MA-005706 | | |

*The original geographical source of this specimen (probably transported to the Netherlands by whalers) is uncertain so calibration with this delta R is an approximation.

**These samples were used to calculate delta R

**Supplementary Table S2.** Confidence scoring of the studied peptide markers for taxonomic assignments.

| **Marker** | **Weight** | **Reason** |
|---|---|---|
| COL1A1 1170–1213 | 3 | Consistently recovered and species-specific |
| Prothrombin | 2 | Consistently recovered and species-specific if known to be Balaenidae |
| COL1A2 262–280 | 2 | Consistently recovered and species-specific if known to be Balaenidae |
| Serotransferrin-like | 1 | Rarely recovered and species-specific if known to be Balaenidae |
| COL1A1 918–957 | 0 | Non-diagnostic due to overlapping polymorphisms |

**Supplementary Table S3.** Summary of specimen type, prior taxonomic identification, peptide-marker detection, and LC-MS/MS-based taxonomic assignment.

| Specimen | Taxonomic Identification (prior to analysis) | Type of Sample | Described in text | Time Period | Collagen peptide markers (Dekker et al. 2024) | Prothrombin | MaxQuant | pFind | COL1A1 918-957 | COL1A1 1170-1213 | COL1A2 262-280 | Prothrombin | Serotransferrin-like | LC-MS/MS taxonomic identity | Markers used |
|---|---|---|---|---|---|---|---|---|---|---|---|---|---|---|---|
| FOC3429 | NARW | Archaeological control | Historical control | Historical | TRUE | TRUE | TRUE | TRUE | Balaenid (both variants) | Not detected | NARW (P variant only) | NARW (E variant only) | Not detected | NARW | Prothombin and COL1A2 262-280 |
| FOC3489 | Bowhead | Archaeological control | Historical control | Historical | TRUE | TRUE | TRUE | TRUE | Bowhead (S variant only) | Bowhead (A variant only) | Bowhead whale (S variant only) | Bowhead whale (D variant only) | Not detected | Bowhead | Prothombin, COL1A1 1170-1213 and COL1A2 262-280 |
| FOC3800 | Bowhead | Red Bay - control for C14 | Historical control | Historical | TRUE | TRUE | TRUE | TRUE | Bowhead (S variant only) | Bowhead (A variant only) | Bowhead whale (S variant only) | Bowhead whale (D variant only) | Not detected | Bowhead | Prothombin, COL1A1 1170-1213 and COL1A2 262-280 |
| FOC3802 | Bowhead | Red Bay - control for C14 | Historical control | Historical | TRUE | TRUE | TRUE | TRUE | Bowhead (S variant only) | Bowhead (A variant only) | Bowhead whale (S variant only) | Bowhead whale (D variant only) | Bowhead whale (R variant only) | Bowhead | Prothombin, Serotransferrin, COL1A1 1170-1213 and COL1A2 262-280 |
| FOC3803 | Bowhead | Red Bay - control for C14 | Historical control | Historical | TRUE | FALSE | TRUE | TRUE | Bowhead (S variant only) | Bowhead (A variant only) | Bowhead whale (S variant only) | Not detected | Not detected | Bowhead | COL1A1 1170-1213 and COL1A2 262-280 |
| FOC3804 | Bowhead | Red Bay - control for C14 | Historical control | Historical | TRUE | FALSE | TRUE | TRUE | Bowhead (S variant only) | Bowhead (A variant only) | Bowhead whale (S variant only) | Not detected | Not detected | Bowhead | COL1A1 1170-1213 and COL1A2 262-280 |
| FOC3820 | Bowhead | Red Bay - control for C14 | Historical control | Historical | TRUE | TRUE | TRUE | TRUE | Bowhead (S variant only) | Bowhead (A variant only) | Bowhead whale (S variant only) | Bowhead whale (D variant only) | Bowhead whale (R variant only) | Bowhead | Prothombin, Serotransferrin, COL1A1 1170-1213 and COL1A2 262-280 |
| FOC3896 | NARW | Archaeological control | Historical control | Historical | TRUE | TRUE | TRUE | TRUE | NARW (A variant only) | Not detected | NARW (P variant only) | NARW (E variant only) | Not detected | NARW | Prothombin and COL1A2 262-280 |
| FOC4050 | Unidentified Balaenidae | Case study | Case study | Case | TRUE | TRUE | TRUE | TRUE | Bowhead (S variant only) | Bowhead (A variant only) | Bowhead whale (S variant only) | Bowhead whale (D variant only) | Not detected | Bowhead | Prothombin, COL1A1 1170-1213 and COL1A2 262-280 |
| FOC4051 | Unidentified Balaenidae | Case study | Case study | Case | TRUE | TRUE | TRUE | TRUE | Bowhead (S variant only) | Bowhead (A variant only) | Bowhead whale (S variant only) | Bowhead whale (D variant only) | Not detected | Bowhead | Prothombin, COL1A1 1170-1213 and COL1A2 262-280 |
| FOC4127 | Unidentified Balaenidae | Case study | Case study | Case | TRUE | TRUE | TRUE | FALSE | Bowhead (S variant only) | Bowhead (A variant only) | Bowhead whale (S variant only) | Bowhead whale (D variant only) | Not detected | Bowhead | Prothombin, COL1A1 1170-1213 and COL1A2 262-280 |
| FOC4192 | NARW | Historical control | Historical control | Historical | TRUE | FALSE | TRUE | FALSE | Balaenid (both variants) | NARW (V variant only) | NARW (P variant only) | Not detected | Not detected | NARW | COL1A1 1170-1213 and COL1A2 262-280 |
| FOC4195 | Unidentified Balaenidae | Case study | Case study | Case | TRUE | TRUE | TRUE | TRUE | NARW (A variant only) | NARW (V variant only) | NARW (P variant only) | NARW (E variant only) | Not detected | NARW | Prothombin, COL1A1 1170-1213 and COL1A2 262-280 |
| FOC4212 | Unidentified Balaenidae | Case study | Case study | Case | TRUE | TRUE | TRUE | TRUE | Bowhead (S variant only) | Bowhead (A variant only) | Bowhead whale (S variant only) | Bowhead whale (D variant only) | Not detected | Bowhead | Prothombin, COL1A1 1170-1213 and COL1A2 262-280 |
| FOC4232 | Unidentified Balaenidae | Case study | Case study | Case | TRUE | TRUE | TRUE | TRUE | Bowhead (S variant only) | Bowhead (A variant only) | Bowhead whale (S variant only) | Bowhead whale (D variant only) | Not detected | Bowhead | Prothombin, COL1A1 1170-1213 and COL1A2 262-280 |
| FOC4238 | Unidentified Balaenidae | Case study | Case study | Case | TRUE | FALSE | TRUE | FALSE | Not detected | NARW (V variant only) | NARW (P variant only) | Not detected | Not detected | NARW | COL1A1 1170-1213 and COL1A2 262-280 |
| FOC4271 | Unidentified Balaenidae | Case study | Case study | Case | TRUE | TRUE | TRUE | TRUE | Bowhead (S variant only) | Bowhead (A variant only) | Bowhead whale (S variant only) | Bowhead whale (D variant only) | Not detected | Bowhead | Prothombin, COL1A1 1170-1213 and COL1A2 262-280 |
| FOC4273 | Unidentified Balaenidae | Case study | Case study | Case | TRUE | TRUE | TRUE | TRUE | Bowhead (S variant only) | Bowhead (A variant only) | Bowhead whale (S variant only) | Bowhead whale (D variant only) | Not detected | Bowhead | Prothombin, COL1A1 1170-1213 and COL1A2 262-280 |
| FOC4274 | Unidentified Balaenidae | Case study | Case study | Case | TRUE | TRUE | TRUE | TRUE | Bowhead (S variant only) | Bowhead (A variant only) | Bowhead whale (S variant only) | Bowhead whale (D variant only) | Not detected | Bowhead | Prothombin, COL1A1 1170-1213 and COL1A2 262-280 |
| FOC4275 | Unidentified Balaenidae | Case study | Case study | Case | TRUE | TRUE | TRUE | TRUE | Bowhead (S variant only) | Bowhead (A variant only) | Bowhead whale (S variant only) | Bowhead whale (D variant only) | Not detected | Bowhead | Prothombin, COL1A1 1170-1213 and COL1A2 262-280 |
| FOC4277 | Unidentified Balaenidae | Case study | Case study | Case | TRUE | TRUE | TRUE | TRUE | Bowhead (S variant only) | Bowhead (A variant only) | Bowhead whale (S variant only) | Bowhead whale (D variant only) | Not detected | Bowhead | Prothombin, COL1A1 1170-1213 and COL1A2 262-280 |
| FOC4282 | Unidentified Balaenidae | Case study | Case study | Case | TRUE | TRUE | TRUE | TRUE | Bowhead (S variant only) | Bowhead (A variant only) | Bowhead whale (S variant only) | Bowhead whale (D variant only) | Not detected | Bowhead | Prothombin, COL1A1 1170-1213 and COL1A2 262-280 |
| FOC4291 | Unidentified Balaenidae | Case study | Case study | Case | TRUE | FALSE | TRUE | FALSE | Not detected | Bowhead (A variant only) | Not detected | Not detected | Not detected | Bowhead | COL1A1 1170-1213 only |
| FOC4314 | NARW | Modern control | Modern control | Modern | TRUE | TRUE | TRUE | FALSE | Balaenid (both variants) | NARW (V variant only) | NARW (P variant only) | NARW (E variant only) | Not detected | NARW | Prothombin, COL1A1 1170-1213 and COL1A2 262-280 |
| FOC4319 | NARW | Modern control | Modern control | Modern | TRUE | TRUE | TRUE | TRUE | Balaenid (both variants) | NARW (V variant only) | NARW (P variant only) | NARW (E variant only) | NARW (K variant only) | NARW | Prothombin, Serotransferrin, COL1A1 1170-1213 and COL1A2 262-280 |
| FOC4323 | NARW | Modern control | Modern control | Modern | TRUE | TRUE | TRUE | FALSE | NARW (A variant only) | NARW (V variant only) | NARW (P variant only) | NARW (E variant only) | Not detected | NARW | Prothombin, COL1A1 1170-1213 and COL1A2 262-280 |
| FOC4326 | Bowhead | Modern control | Modern control | Modern | TRUE | TRUE | TRUE | TRUE | Bowhead (S variant only) | Bowhead (A variant only) | Bowhead whale (S variant only) | Bowhead whale (D variant only) | Not detected | Bowhead | Prothombin, COL1A1 1170-1213 and COL1A2 262-280 |

**Supplementary Table S4.** Peptide marker recovery rates.

| Marker | Overall recovered / total | Overall recovery rate | Overall Wilson 95% CI | Overall 95% rarefaction range (n = 10) | Non-modern recovered / total* | Non-modern recovery rate* | Non-modern Wilson 95% CI | Non-modern 95% rarefaction range (n = 10) | Modern recovered / total | Modern recovery rate | Modern Wilson 95% CI |
|---|---|---|---|---|---|---|---|---|---|---|---|
| COL1A1 918-957 | 25 / 27 | 92.6% | 76.6–97.9% | 80–100% | 21 / 23 | 91.3% | 73.2–97.6% | 80–100% | 4 / 4 | 100.0% | 51.0–100.0 % |
| COL1A1 1170-1213 | 25 / 27 | 92.6% | 76.6–97.9% | 80–100% | 21 / 23 | 91.3% | 73.2–97.6% | 80–100% | 4 / 4 | 100.0% | 51.0–100.0 % |
| COL1A2 262-280 | 26 / 27 | 96.3% | 81.7–99.3% | 90–100% | 22 / 23 | 95.70 % | 79.0–99.2% | 90–100% | 4 / 4 | 100.0% | 51.0–100.0 % |
| Prothrombin | 22 / 27 | 81.5% | 63.3–91.8% | 60–100% | 18 / 23 | 78.3% | 58.1–90.3% | 60–100% | 4 / 4 | 100.0% | 51.0–100.0 % |
| Serotransferrin-like | 3 / 27 | 11.1% | 3.9–28.1% | 0–30% | 2 / 23 | 8.7% | 2.4–26.8% | 0–20% | 1 / 4 | 25.0% | 4.6–69.9% |
| *Non-modern recovery rates exclude the four modern control samples | | | | | | | | | | | |

**Supplementary Table S5.** Diagnostic peptide marker recovery rates.

| Marker | Species-specific diagnostic recovery across controls | Bowhead-specific recovery in *B. mysticetus* controls | NARW-specific recovery in *E. glacialis* controls | Bowhead-specific recovery in modern *B. mysticetus* controls | NARW-specific recovery in modern *E. glacialis* controls | Bowhead-specific recovery in historical *B. mysticetus* controls | NARW-specific recovery in historical *E. glacialis* controls |
|---|---|---|---|---|---|---|---|
| COL1A1 918–957 | 0 / 13 = 00.0% | 0 / 7 = 00.0% | 0 / 6 = 00.0% | 0 / 1 = 00.0% | 0 / 3 = 0.0% | 0 / 6 = 0.0% | 0 / 3 = 0.0% |
| COL1A1 1170–1213 | 11 / 13 = 84.6% | 7 / 7 = 100.0% | 4 / 6 = 66.7% | 1 / 1 = 100.0% | 3 / 3 = 100.0% | 6 / 6 = 100.0% | 1 / 3 = 33.3% |
| COL1A2 262–280 | 13 / 13 = 100.0% | 7 / 7 = 100.0% | 6 / 6 = 100.0% | 1 / 1 = 100.0% | 3 / 3 = 100.0% | 6 / 6 = 100.0% | 3 / 3 = 100.0% |
| Prothrombin | 10 / 13 = 76.9% | 5 / 7 = 71.4% | 5 / 6 = 83.3% | 1 / 1 = 100.0% | 3 / 3 = 100.0% | 4 / 6 = 66.7% | 2 / 3 = 66.7% |
| Serotransferrin-like | 3 / 13 = 23.1% | 2 / 7 = 28.6% | 1 / 6 = 16.6% | 0 / 3 = 0.0% | 1 / 3 = 33.3% | 2 / 6 = 33.3% | 0 / 3 = 0.0% |

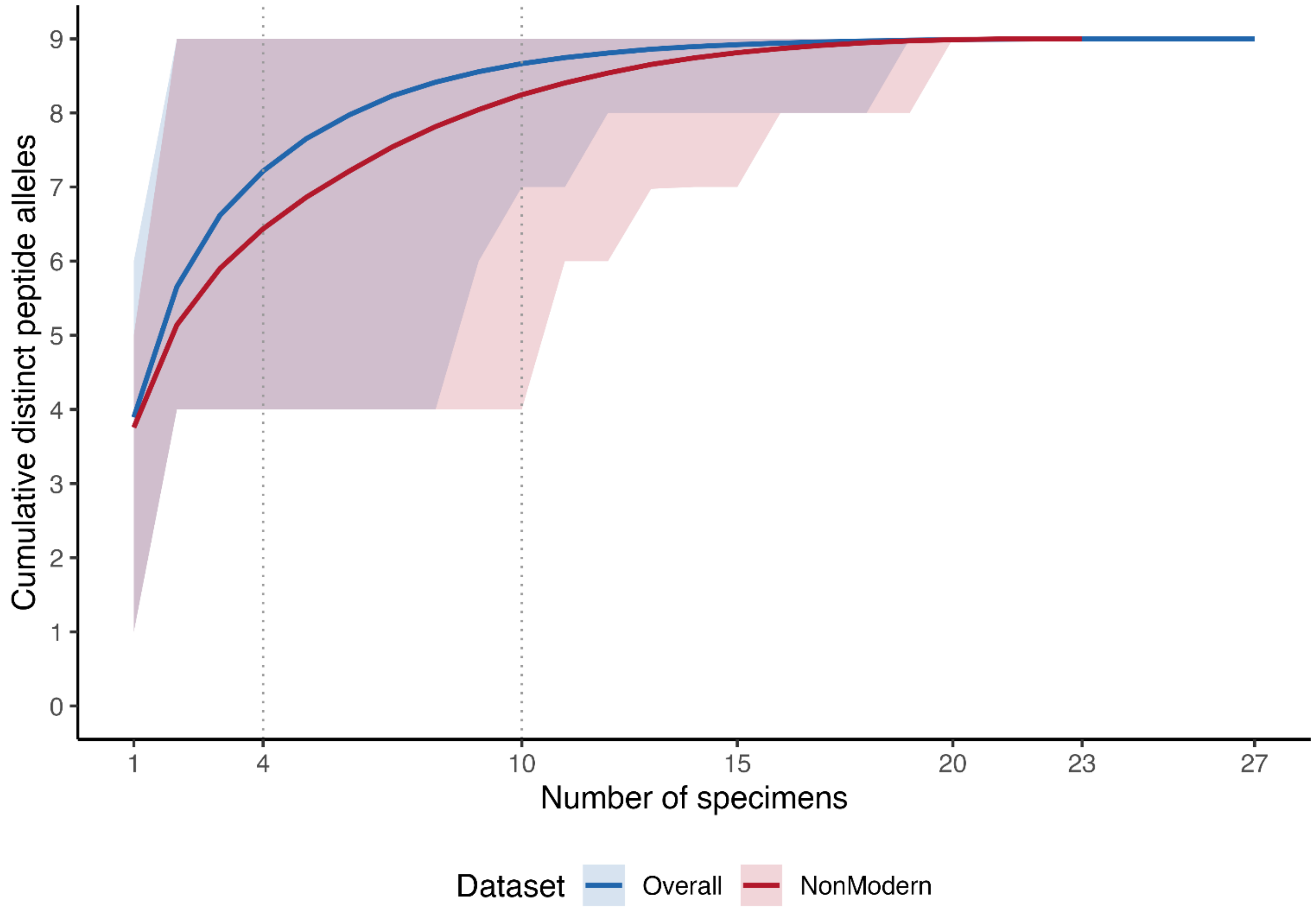


**Supplementary Figure S1.** Sample-based rarefaction of peptide variant accumulation. Curves show the mean cumulative number of distinct peptide variants recovered with increasing specimen numbers for the overall (n=27; blue) and historical (n=23; red) datasets. Rarefaction was performed using 10,000 random permutations of specimens without replacement. Shaded areas represent 95% rarefaction ranges. Vertical dotted lines indicate sampling depths at four and ten specimens.
